# Sub-clinical triiodothyronine levels predict health, demographic, and socioeconomic outcomes

**DOI:** 10.1101/2023.03.09.531775

**Authors:** Ralph I. Lawton, Bernardo L. Sabatini, Daniel R. Hochbaum

## Abstract

The Hypothalamic-Pituitary-Thyroid (HPT) axis is fundamental to human biology, exerting central control over energy expenditure, metabolic rate, and body temperature. However, the consequences of “normal” physiologic HPT-axis variation in non-clinical populations are poorly understood. Using nationally-representative data from the 2007-2012 NHANES, we explore relationships with demographics, mortality, and socio-economic factors. We find much larger variation across age in free T3 than other HPT-axis hormones. T3 and T4 have opposite effects on mortality: free T3 is inversely related and free T4 is positively related with likelihood of death. Free T3 and household income are negatively related, particularly at lower incomes. Finally, free T3 among older adults is associated with labor both on the extensive margin (unemployment) and intensive margin (hours worked). Physiologic TSH/T4 explain only 1% of T3 variation, and neither are appreciably correlated to socio-economic outcomes. Taken together, our data suggest an unappreciated complexity and non-linearity of the HPT-axis signaling cascade broadly such that TSH and T4 may not be accurate surrogates of free T3. Furthermore, we find that sub-clinical variation in the HPT-axis effector hormone T3 is an important and overlooked factor linking socio-economic forces, human biology, and aging.

## Introduction

The Hypothalamic-Pituitary-Thyroid (HPT) axis exerts central control over core components of human biology, including energy expenditure, metabolic rate, and body temperature regulation. Abnormal levels of thyroid hormone (either hypo- or hyper-thyroid) result in metabolic dysregulation and myriad psychiatric disease symptoms (1–7). Despite the clinical focus on hyper- and hypo-thyroidism, most individuals’ thyroid hormones have ‘set-points’ that vary sub-clinically, and laboratory reference ranges are large and therefore insensitive to normal variation in thyroid function (8–10). The prominence of both metabolic and psychological phenotypes in thyroid disease-states suggests that the HPT axis could govern aspects of health and human behavior within normal, sub-clinical variation as well. Further, given the declines in HPT-axis function and metabolism with aging, these sub-clinical set-points may be important mediators of well-being among older populations (11–13). However, the role of normal physiological variation of the HPT-axis in human behavior, socio-economic forces, and population health remains poorly understood, particularly with respect to the HPT-axis effector hormone T3 (8).

The canonical HPT-axis signaling cascade begins with the hypothalamus stimulating the pituitary gland to release TSH (thyroid stimulating hormone), TSH stimulates the thyroid gland to secrete primarily T4 (thyroxine) and some T3 (triiodothyronine). The vast majority of T3 is produced in the periphery of the body, where T4 is converted into T3 by deiodinases DIO1 and DIO2. T3 is considered to be the active hormone, with ∽10-to 30-fold higher affinity for thyroid receptors than the prohormone T4 (14). Changes in the levels of T3 alter function of tissues throughout the body (15). Clinically, hypothyroidism tends to lead to lethargy, weight gain, hypothermia, and depression, whereas hyperthyroidism tends to lead to weight loss, hyperthermia, agitation, restlessness and hyperactivity, sleep disruption, and other manic-like symptoms. Given the centrality of thyroid function to bodily homeostasis and energy levels, and the emergence of altered behaviors when dysregulated, one would expect socio-economic forces to both shape and be shaped significantly by thyroid function. Yet little attention has been given to sub-clinical variation in HPT-axis function.

Understanding the role of the HPT-axis in aging and population health is complicated by mixed evidence from studies of the HPT-axis and mortality. Although low T3 and T4 levels have been linked to adverse mortality outcomes in hospitalized or cardiac patients, the role of HPT-axis hormones in non-clinical populations is less clear (16, 17). Evidence from euthyroid patients visiting clinics in South Korea found both free T3 and free T4 to be protective against mortality, whereas other studies of euthyroid patients in Europe have found free T4 to be negatively related to mortality (18–20). Interventions with levothyroxine (synthetic T4) have not been protective against mortality in older adults but may be beneficial in younger adults (21).

Here, we use nationally-representative data from the USA to study the relationships between HPT-axis function in the general population across the physiological range with demographics, core functions of the HPT-axis, socio-economic forces, and mortality. We use simultaneous measures of TSH, free T4, and free T3, enabling thorough assessment of HPT-axis hormones, and correlate their levels with key dimensions of wellbeing, especially as people age.

## Results

We combined data on TSH, free T4, and free T3 from 8,059 adults over age 20 the 2007-2012 survey waves of the National Health and Nutrition Examination Survey (NHANES). Weighted summary statistics are in supplement table S1. To account for potential untreated clinical hypo- and hyper-thyroidism, we drop individuals outside the top and bottom 1% of the sample (n=691). For all regression results, we estimate three models to examine relationships between covariates and thyroid hormones. We specify a base model conditioning on age, demographics, survey wave, smoking, and thyroid-related medication use, and extend this model to include measures of socio-economic status (SES), as well as measures of health that may be related to thyroid function (see Methods). We focused on free T3 and T4 since protein-bound T4 and T3 not thought to be biologically active. Simultaneous measures of TSH, free T4, and free T3 in each survey participant enables localization of specific signaling stages in which the HPT-axis modifies and is shaped by population health and socio-economic forces.

### Demographics and Seasonality

We sought to confirm previous findings relating basic demographics and seasonality to HPT-axis function to indicate the robustness of our chosen dataset. Consistent with evidence from the 2001 NHANES that studied TSH but not free T3 or T4, we find that males and females have statistically indistinguishable TSH levels. We also find small differences in free T4 which are not robust to controls for other health measures (Table 1) (22).^1^ Additionally, we find that males have 0.5 standard deviations (0.17pg/mL) higher free T3 levels (p<0.001), consistent with data from clinical samples (23, 24). Examining differences by race replicates prior findings with respect to TSH (Table 1) (22). Black Americans have 0.4 standard deviations lower TSH and 0.1 standard deviations lower free T3 than non-Hispanic white Americans (p<0.01). Hispanic Americans have 0.1 standard deviation lower TSH, 0.2 standard deviation higher free T3 (p<0.01). and 0.1 standard deviation higher free T4 (p<0.05), though free T4 results are not robust to SES controls. Finally, evidence from mammalian studies implicates seasonality in thyroid function, as do recent meta-analyses of human thyroid studies, attributed largely to the role of thyroid function in thermoregulation (25–27). Consistent with these studies, we find evidence for lower free T3 during the summer (Table 1, **Figure 1**). Therefore, the 2007-2012 waves of the NHANES recapitulate canonical findings of HPT-axis function in a nationally-representative sample.

**Figure 1.**
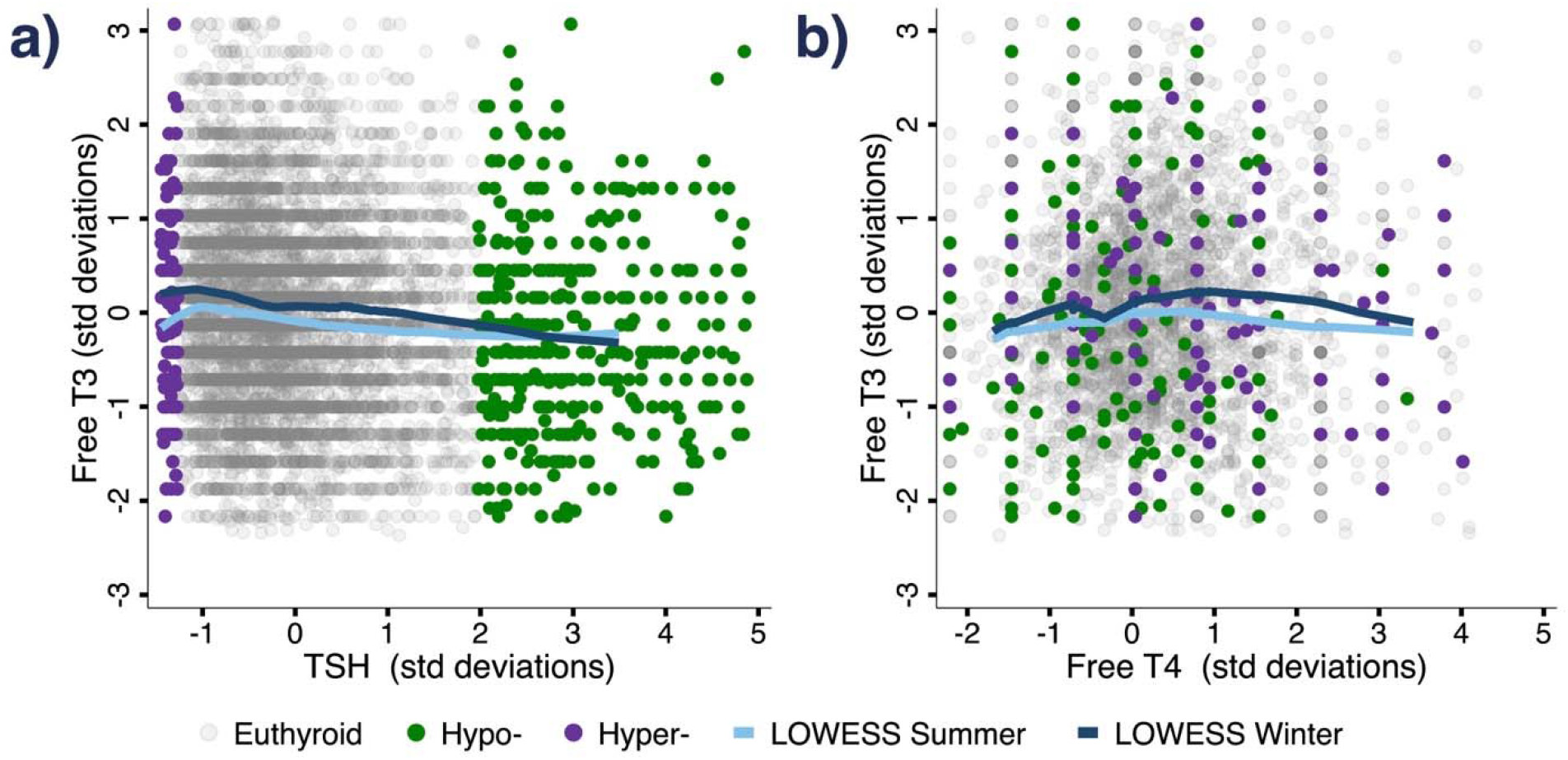
Inter-relationships between TSH, free T4, and free T3 in standard deviations. a) Scatter plot showing TSH and free T3 for the same individual, in terms of standard deviations of the adult population distribution. b) Scatter plot showing free T4 and free T3 for the same individual, in terms of standard deviations of the adult population distribution. Euthyroid, hypo-, and hyper-thyroid individuals in both panels classified by TSH (hypo: TSH>4.1 mIU/L, hyper: TSH<0.4 mIU/L). Non-parametric LOWESS estimates shown stratifying by summer measurement (May 1-Oct. 31). **Supplement Figures S1 and S2** display parallel results in non-standardized units, and stratifying overt and sub-clinical hypothyroidism.

### Relationships between TSH, T4, and T3

Although TSH, T4, and T3 are considered a canonical chain of HPT-axis hormones that are tightly coupled, no population-representative studies exist that measure all three simultaneously within individuals to validate this relationship. Evidence from clinical samples suggests that HPT-axis hormones are not strongly predictive of one another, as free T4 and even TSH can be unreliable measures of sub-clinical but symptomatic hypothyroidism (13, 28, 29). Further, 5-10% of hypothyroid individuals treated with levothyroxine (LT4) that successfully normalize TSH levels remain symptomatic, suggesting that normal TSH and T4 levels can be insufficient to restore normal T3-mediated physiology (30).

We therefore directly compared TSH, free T4, and free T3 within individuals to evaluate the correlation between these HPT-axis molecules. We find that the relationships between TSH (**Figure 1a, Supplemental Fig 1a**) and free T4 (**Figure 1b, Supplemental Fig 1b**) with free T3 are weak, explaining 0.4% and 0.8% of the variation in free T3 respectively, and 1% combined (Supplement Table S2). Even clinical thresholds for hyper-, hypo-, and subclinical hypo-thyroidism based on TSH and free T4, poorly stratify free T3 levels (**Supplemental Fig 2**). Hypothyroid adults, classified by classified by high TSH (TSH > 4.1mIU/L, 1.98SD, Garber et al.) and low free T4 (T4 < 0.7 ng/dL, 0.86SD, see methods) are only 1/3 of a standard deviation lower in their free T3 levels conditional on our base model (6). Sub-clinical hypothyroidism, classified by high TSH and ‘normal’ T4 (>0.7ng/dL), is even more poorly stratified, with low ability to stratify free T3 levels (**Supplemental Fig 2a-b**), and no ability to stratify free T3 conditional on age, demographics, smoking, and seasonality (Supplement Table S2) (6). Even hyperthyroidism (TSH<0.4mIU/L, Garber et al.) does poorly, with functionally no relationship to free T3 (6).^2^ Therefore, free T3 levels cannot be reliably predicted by TSH and free T4 levels, and direct measurement of free T3 is likely vital to properly stratify the effects of HPT-axis variation on clinical and non-clinical populations.

### Aging and mortality

Age is one of the strongest predictors of clinical hypothyroidism, and age-related HPT-axis function has been implicated in the decline of metabolic rate and increase in adiposity as adults age (22, 31, 32). However, prior work has only focused on TSH and free T4 levels, which are commonly used in the clinical diagnosis of thyroid disease. Non-parametric models stratified by sex show the relationships between all three standardized HPT-axis hormones and age (**Figure 2a**). Free T3 exhibits substantially more variation across age than either free T4 or TSH, with over twice the variation between the oldest and the youngest than TSH. Free T3 declines with age, consistent with its role as the active effector molecule of the HPT-axis, whose function declines with age. While total T3 follows similar patterns to free T3, the relationships with age are attenuated (**Supplement Figure S3**), likely due to unaccounted variability of T3 binding proteins. TSH levels steadily increase over time, whereas the free T4 pattern is nonlinear, declining until approximately age 50 then increasing. Given that hypothyroidism is typically diagnosed clinically as high TSH levels and low T4, the simultaneous increases later in life in both TSH and free T4 could potentially confound diagnosis of clinical and sub-clinical hypothyroidism (6). Further, the diverging levels of T4 and T3 with age suggests altered conversion of T4 to T3, potentially mediated by age-related degradation of deiodinase function.

**Figure 2.**
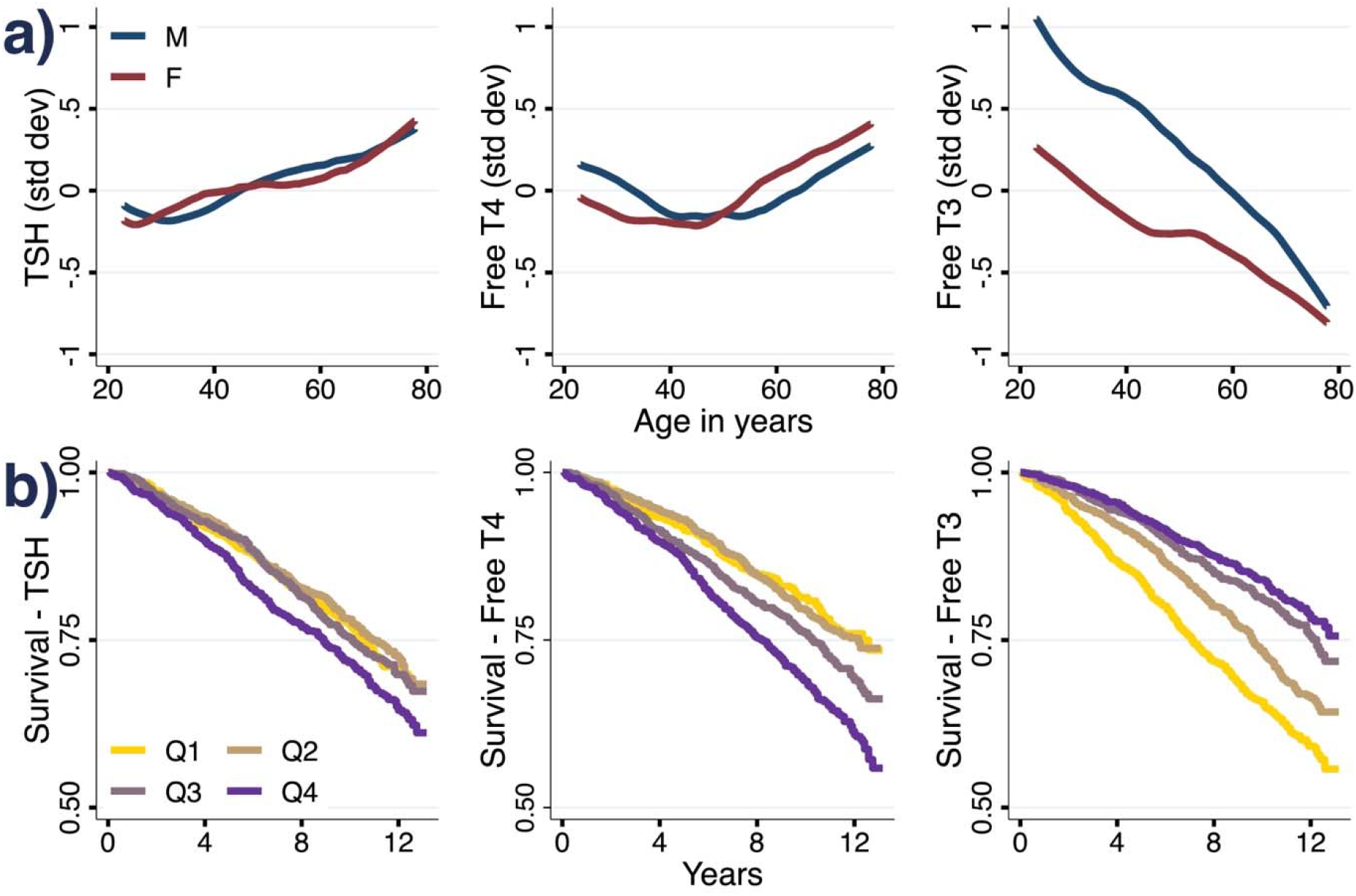
Thyroid hormones, age, and mortality. **A)** weighted non-parametric LOWESS estimates of the relationships between thyroid hormones and age among adults, stratified by sex. **B)** weighted survival curves since measurement date for each quartile of TSH, free T4, and free T3, among adults over age 50 (Q1: 0-25%, Q2: 25-50%, Q3: 50-75%, Q4: 75-100% within each hormone’s distribution).^3^

Given the prominent variation with age, we next examined if HPT-axis hormones are predictive of mortality. Using mortality data through the end of 2019 linked to the NHANES, we use Cox proportional-hazards models evaluate the relationships between HPT-axis hormones and mortality in a non-clinical population. Kaplan Meier curves separated by quartile of each hormone are shown in **Figure 2b**. We focused on adults over 50 (n = 3,953), among whom there were 1,086 deaths in this time period. TSH is not linked to mortality, but increasing levels of free T3 and decreasing levels of free T4 are protective from mortality (hazard ratios 0.87 and 1.19, respectively, p<0.01), and retain their coefficients conditional on socioeconomic status (SES) and important markers of health including measures of adiposity like waist circumference (Supplement Table S3). Higher T3 quartiles have slightly higher BMI than lower ones (Q4: 29.46, Q1: 28.79), but as BMI is linked to higher mortality, these should bias mortality estimates the opposite direction of the T3 relationship. Total T3 has a similar relationship to mortality as free T3, but total T4 is attenuated significantly from free T4’s mortality relationship (**Supplement Figure S3**, Supplement Table S4). As illness can affect thyroid hormone levels and directly affect mortality, we evaluate the potential role of illness by extending our full model to include covariates for flu, colds, or GI illness in the past 30 days, as well as measures of inflammation using C-Reactive Protein from the first two waves of data. Mortality results are robust to these controls, though some statistical power is lost when only using the first two waves (Supplement Table S5).

Notably, these biomarkers are not predictive of mortality simply in the short term but continue to stratify mortality through the long-term endpoint (**Figure 2b**). These results indicate that age-related changes in T3 and T4 levels are important factors related to longevity. Further, because T3 and T4 levels have opposing relationships to mortality (high T3 and low T4 are protective from mortality), the peripheral conversion of T4 to T3 by deiodinases is likely a critical determinant of longevity.

### Socio-Economic Status and Employment

We examined the correlation between social forces such as household resources and thyroid function. Conditional on prior discussed covariates, we find a robust negative relationship between household income and free T3, but no relationship between household income and free T4 (Table 1, **Figure 3**). Although there is a negative relationship between income and TSH (**Figure 3a**), it is not robust to inclusion of other health factors that may affect the HPT-axis, nor to an extended model that controls for free T3; however, the free T3 relationship is robust to controlling TSH. The negative relationship between free T3 and socio-economic status (SES) persists with respect to alternative measures of SES including household income per capita, and income-to-poverty ratios that account for household size, state, and year specific poverty thresholds (Supplement Table S6). Similar models for Total T3 find similar, but less-precisely estimated relationships between income and total T3 than free T3 (**Supplement Figure S7**, Supplement Table S8).

**Figure 3.**
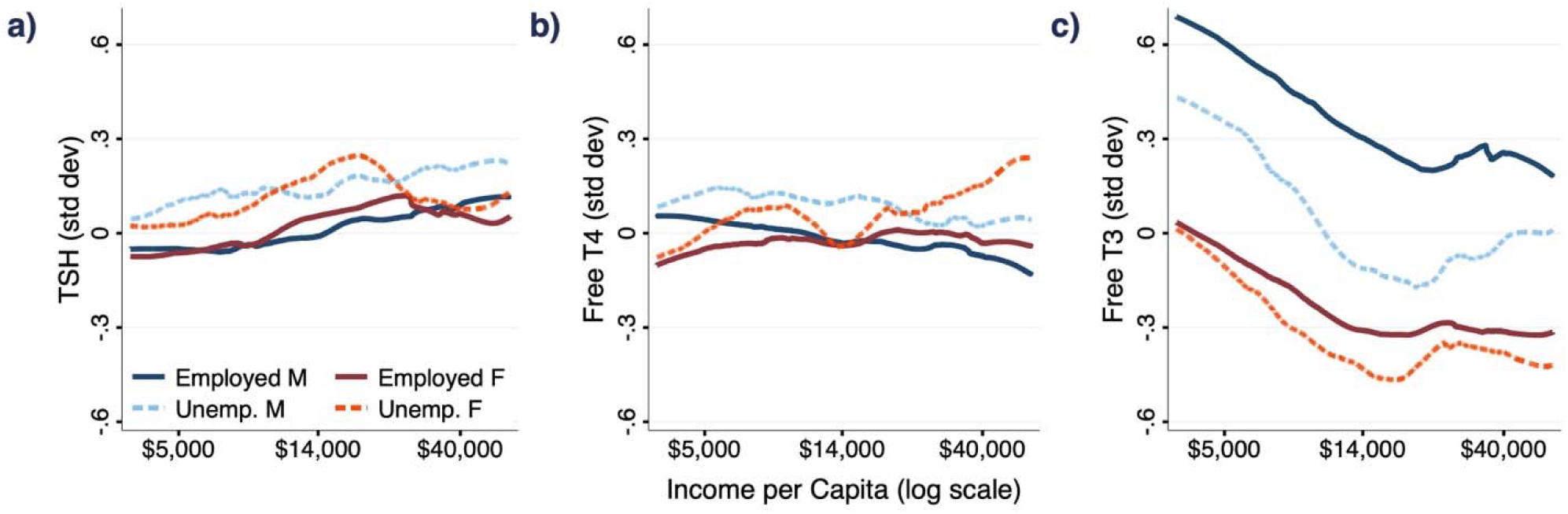
Thyroid hormones, household income, and unemployment. Weighted non-parametric LOWESS estimates of the relationship between thyroid hormones and the natural logarithm of real (2007 dollars) household income per capita are displayed, stratified by sex and employment status at the time of measurement. Data from 2007-2012 NHANES.

The negative relationship between free T3 and household resources is stronger at lower income levels (**Figure 3c**). We estimate the location of a ‘kink’ in the relationship between income and free T3 (**Figure 3c)**, and conduct statistical tests for the significance of a threshold effect relative to a linear model (Supplement B1) (33). Our analysis identifies the threshold at $22,735 dollars per capita and finds the ‘kink’ model to be a significantly better fit than the linear model (p<0.001). $22,735 is approximately half of average income per capita in 2007, suggesting an interaction between stressors associated with lower-income households and T3 (34).

Although this observed correlation cannot be interpreted causally, the relationship is nearly identical for employed and unemployed adults across both genders (**Figure 3c**), suggesting that potential links between T3 and earnings in the job market are not driving the income-T3 correlation directly, but rather implicating a household-level force related to resources. The effects are meaningfully large in magnitude, similar to or larger than for other biomarkers with important income-gradients like cholesterol, % HbA1C, blood pressure, and C-reactive protein (Supplement Table S7) (35, 36).

Although household resources and free T3 are inversely related, **Figure 3** shows free T3, but not free T4 or TSH, is lower among unemployed men. We used ordinary least squares (OLS) and logit models to explore potential relationships between free T3 and employment status among men. Since age is a key mediator of both employment status and free T3, we specify a fully-interacted model of our full specification’s covariates interacted with age, that can account for co-variation in the effects of free T3 and employment status as individuals age (see Methods). We find evidence that at older ages, free T3 is a key mediator of employment status (**Table 2**; see **Supplemental Table S9** for similar results using total T3). Logit models find that for each decade increase in age, a 1 SD increase in T3 is linked to an improvement in the odds of employment by 1.18. Non-parametric explorations bear this effect out: among adults under 40, there is no apparent relationship between free T3 and employment status; however, among adults over 40, there is a clear positive relationship, which is particularly salient at lower levels of free T3 (**Supplement Figure S4**).

These effects are apparent on both the extensive and intensive margin. Regression models estimating the effect of free T3 on the number of hours worked overall and conditional on employment show increases not only in labor market participation but also in the numbers of hours worked (**Table 2, Figures S5, S6**). These relationships with hours worked are strongest among those employed in ‘high-activity’ jobs, classified as manual or service jobs, among whom there is not just an effect of free T3 at older ages but a large effect of free T3 irrespective of age on the number of hours worked (37). The association between employment and free T3 is consistent with the effects of increased free T3 on energy expenditure— effects that are observed clinically in the context of hyperthyroidism with increased metabolism in addition to manic and hyperactive psychiatric symptoms.

Notably, across all models, we can explain a much larger (∽2-5x) proportion of variation in free T3 than in free T4 or TSH (**Table 1**). Whereas we can explain 24% of the variation in free T3, we only can explain 5-10% of the variation in free T4 or TSH with the same models.

Collectively, these results demonstrate that free T3 is the sole HPT-axis molecule predictive of socio-economic outcomes. Given that maintenance of free T3 levels is held in equilibrium by both production of T4 and deiodinase activity converting T4 to T3, our results suggest that peripheral conversion of T4 to T3 responds in important ways to demographic and socio-economic variables of interest.

## Discussion

The HPT-axis is central to many facets of human biology, including energy expenditure, metabolic rate, and body temperature regulation. The physiological and behavioral ramifications of hyper- and hypo-thyroidism have been well-studied. However, most individuals exhibit sub-clinical variation in HPT-axis hormone levels (8–11, 31) and the ramifications of ‘normal’ thyroid function in the general population and among non-clinical samples has been minimally-explored (31). In particular, very few studies investigate the correlates of variation in free T3 (11, 31).

In a non-clinical sample representative of the US population, we find that free T3 is much more strongly related to all domains studied— age, sex, seasonality, household income, employment, and mortality— than the other molecules of the HPT-axis. The relative strength of these relationships reinforces free T3’s privileged role as the primary effector of the HPT-axis, and suggests that for population research, and likely for clinical surveillance, increased focus on free T3 as a primary readout of HPT-axis function is warranted.

These findings have particular salience for the wellbeing of aging populations. Although it is known that HPT-axis function declines with age, measured variation in free T3 is much larger than the variation in free T4 or TSH, and only T3 steadily declines with age. Furthermore, free T4 and free T3 diverge at older ages, when free T4 increases and free T3 continually decreases. This may reflect age-related decline in the efficiency of T4 to T3 conversion by deiodinases (12, 38–40). Our sample is nationally-representative and covers a wide age spectrum from ages 20-80. Other studies of centenarians suggest the patterns we observe may continue at even older ages (12). Given known declines in HPT-axis function as people age, and the typical use of free T4 and TSH in diagnosis of hypo-thyroidism, diverging patterns in free T4 and free T3 with age have significant implications for the proper diagnosis and HPT-axis assessment at older ages: older adults with symptoms of hypothyroidism may have high free T4 levels despite low free T3.

The divergence between free T3 and free T4 at older ages is paralleled in our mortality findings. Whereas increasing free T3 is protective against mortality, increasing free T4 is linked to higher levels of mortality, suggesting that the age-related decline in T4 to T3 conversion, and the role of deiodinases in this process, have important impacts on longevity. These results parallel similar findings on free T3, T4, and deiodinase function in hospitalized and frail patients, but suggest further functional importance at sub-clinical levels in the general population (41). Poor deiodinase function at older ages may also reconcile why levothyroxine (synthetic L-T4) therapy does not improve mortality outcomes among the oldest patients with sub-clinical hypothyroidism, but may be helpful at younger ages (21). Recent findings have suggested a potentially important clinical role for free T3 measurement, as well as benefits of combination liothyronine and levothyroxine therapy in highly symptomatic patients. (42, 43)

We find a robust negative relationship between household resources and free T3, to our knowledge the first evidence of social modification of thyroid hormone profiles in the general population. We do not see this pattern for either free T4 or TSH. This evidence is consistent across employed and unemployed household members of both sexes, suggesting that direct effects of income generation are not driving the observed pattern. The mechanism for this result is not obvious, but could be related to household resource-related stress (34). The relationship is primarily present at lower levels of income, suggesting a threshold at which the marginal effects of income on free T3 levels diminish.

Stratifying our income relationships by employment status does not reveal differential effects. Instead, doing so emphasizes a significant drop in free T3 levels among unemployed versus employed males. This may have important implications for well-being, especially among older males. We find that at ages 40 and older, free T3 significantly affects the likelihood of being employed. This relationship is apparent on both the extensive margin and the intensive margin, with similar patterns in employment status as well as hours worked among those who are employed. Our findings may suggest that normal variation (non-pathological) in levels of T3 influences individuals’ willingness to engage and be active, particularly at older ages. These relationships are consistent with T3’s role in governing energy expenditure, the prominence of hyperactive psychiatric symptoms in hyperthyroidism, and recent evidence from animal models on the role of free T3 on exploratory activity and behavior, as well as evidence suggesting treating overt clinical hypothyroidism leads to improved employment outcomes (44, 45).

These results underscore the potential importance of measuring and understanding free T3 as the primary effector molecule of the HPT-axis. It is important to note that the more-typically measured biomarkers for thyroid function (T4 and TSH) are poorly linked to free T3 levels. We document significant non-linearity in the relationships between TSH, T4, and T3, and find less than 1% of the variation in free T3 is explainable with these other measures.

The evidence presented here on the central role of free T3 also gives insight into the potential mechanisms involved. Free T3 is much more strongly related than other HPT measures to core HPT-axis dimensions like seasonality, sex, and age, as well as socio-economic status and employment. Furthermore, T3 diverges from free T4 with respect to age and mortality. Thus, our findings suggest a vital importance of the efficiency of conversion between T4 and T3 both in determining T3 levels and their relationship to socio-economic factors. Given the known role for deiodinases in these processes, further studies of the sub-clinical regulation of deiodinase activity by social forces and aging could elucidate critical biology governing these effects, while new therapeutics targeting deiodinase activity may be more beneficial for a significant cohort of older individuals currently treated with levothyroxine.

## Data and Methods

### Data

We combine data from the 2007-2008, 2009-2010, and 2011-2012 waves of the NHANES, a nationally representative sample of the United States population. These are the only recent waves of the NHANES study that measure free T3, free T4, and TSH.^4^ While all adults in the 2007-2008 NHANES were eligible for HPT-axis measures, only sub-samples of the 2009-2010 and 2011-2012 waves were selected to have HPT-axis measures collected (46). We focus on adults over age 20 at the time of survey, and exclude women who were pregnant at the time of assessment. To account for potential untreated clinical hyper and hypothyroidism, we drop individuals outside the top and bottom 1% of the sample.^5^

Our core outcomes are the key hormones of the HPT-axis: TSH, free T4, and free T3. Canonically, the hypothalamus stimulates the pituitary gland to release TSH, TSH stimulates the thyroid gland to secrete primarily T4 and some T3, and the vast majority of T3 is produced in the periphery of the body, where T4 (considered a prohormone with limited activity) is converted into the active hormone T3 by deiodinase enzymes (5). T3 and T4 both circulate in free (active) and protein-bound forms (inactive). Across all NHANES waves used, thyroid hormone concentrations were assessed using a Beckman Coulter Access 2 immunoassay, validation documentation publicly available from the NCHS (46).

Linked mortality information through 2019 is sourced from the NCHS public-use linked mortality dataset. Since mortality among individuals under 50 is very rare, we focus our mortality analysis on adults over age 50 at the time of measurement.

Age is reported by the respondent. In this data, it is top-coded at age 80. Race and ethnicity data, education data, household size, smoking behavior, and country of origin is based on self-report. ‘Summer’ measurement is defined as measurement in the six-month period between May 1 and October 31. Employment status is based upon whether respondents reported they were ‘working at a job or business’ in the last week. Income is assessed at the household level, and is adjusted across waves for inflation to 2007 dollars.

Where used, clinical cutoffs for hypothyroidism, sub-clinical hypothyroidism, and hyperthyroidism were defined as follows: hypothyroidism (TSH>4.1 mIU/L, free T4<15^th^ percentile), sub-clinical hypothyroidism (TSH>4.1 mIU/L, free T4>15^th^ percentile), and Hyperthyoidism was defined as (TSH<0.4 mIU/L). Classifications of high and low TSH approximate 2012 guidelines from the AACE/ATA, and typical laboratory reference ranges from recent empirical work (47–49). Percentiles were used for free T4 to conservatively categorize hypothyroidism from sub-clinical hypothyroidism given lack of clear guidelines for sub-clinical hypothyroidism and free T4, as older work like the Colorado Thyroid Disease Prevalence Study or NHANES III used total T4 levels, and very few adults (<0.5%) had free T4 levels over 1.8ng/dL as suggested by Kim et al. (47, 48, 50, 51).

### Statistical Analysis

To account for the complex sampling design of the NHANES, all models were adjusted for sampling weights, clustering, and stratification. Since the 2009-2010 and 2011-2012 waves were 1/3 sub-samples with specific sub-sample weights to represent the full population in those waves, the weights were divided by three, such that within each wave the weights still are population-representative, but that across waves individual observations are similarly weighted. Analyses were conducted using the SVY routine with linearized standard errors in Stata/SE 17.0.

In order to examine associations between the HPT-axis hormones and age, a known driver of thyroid function, we used LOWESS to explore non-parametric relationships (52).

Next, we used a series of OLS models to 1) establish core demographic relationships between age, gender, race, and seasonality of measurement and 2) examine the potential relationships between HPT-axis hormones and socio-economic gradients, focusing on household resources. We specify three models, all of which are conditional on thyroid-related medication use, statin use, and wave. For the first model, we examine relationships between age, gender, race, and seasonality on HPT-axis hormones, conditional on medication use, wave, and smoking behavior. Due to non-linearities in HPT-axis hormones and age, particularly free T4, we model age semi-parametrically with a series of dummy indicators for each decade.^6^ Second, we extend the first model to real household income, college education, household size, and nativity. Finally, we extend the model further to investigate whether our demographic and economic relationships of interest are confounded by common factors that affect the HPT-axis, namely body composition, sleep, iodine levels, nutrition, and HbA1c as a proxy for long-term blood glucose.

In order to study mortality, we apply the same series of models described above. We use Cox proportional hazards models specified on the months that have passed between an interview date and mortality in order to account for varying time between each wave and the 2019 mortality data endpoint.

We also investigate the interplay between the HPT-axis and employment status. Specifying both OLS and logistic regression models conditional on the same covariates described above, we investigate the relationships between HPT-axis hormones and the likelihood an individual is currently employed. Since both employment status and HPT-axis hormones have strong relationships with age, we specify fully-interacted models that allow the effects of HPT-axis hormones on employment to vary with age.

In order to supplement our regression analyses of the relationships between HPT-axis hormones with income and employment, we use LOWESS to examine non-linearities in these relationships (52). With respect to employment, in order to visualize the interaction between age and free T3 with employment, we stratify into three age categories, standardize free T3 within each group, and display the LOWESS models of employment probability. Observing that the non-parametric relationship between income and free-T3 is downward-sloping at lower income levels and functionally flat at higher levels, we follow Hansen (2017) to estimate the location of a potential threshold at which the slopes change, and test for whether the regression fit on either side of the threshold are significantly different from the base linear model (Supplement B1).

## Footnotes

1 While Aoki et al. did not study free T4, our results on free T4 are similar to theirs on total T4. Conditional on age and race, Aoki et al. finds small but statistically significant differences in total T4 by sex. We find similar results in free T4 with our base model, similar to Aoki et al., but conditioning on further health measures renders the estimate non-significant.

2 Excluding individuals on thyroid medications does not affect the relationships between hypothyroid and free T3, but further attenuates relationships between sub-clinical hypothyroid and hyperthyroid classifications.

3 TSH: Q1=0.201-1.15uIU/mL, Q2=1.151-1.745uIU/mL, Q3=1.746-2.58uIU/mL, Q4: 1=2.581-7.405uIU/mL. Free T4: Q1=0.5-0.69ng/dL, Q2=0.7-0.79ng/dL, Q3=0.8-0.89ng/dL, Q4=0.9-1.35ng/dL. Free T3: Q1=2.33-2.8pg/mL,Q2=2.81-3pg/mL, Q3=3.01-3.2pg/mL, Q4=3.21-4.21pg/mL.

4 While we are not fully certain why TSH, and to a lesser extent T4, have been the focus of prior population-based literature, the CDC stopped collecting data on T4 and T3 after 2012, saying in email correspondence that there was no further public-health support for T3 collection.

5 Our results are not sensitive alternative specifications that winsorize at 5 or 10%, nor are they sensitive to dropping all individuals with clinical hypo and hyperthyroidism (Supplement Table S10, S11, S12, S13). Weighted summary statistics are presented in Supplement Table S1.

6 Nonetheless, results are robust to linear instead of semi-parametric controls for age (Supplement Table S14).

