## Supplementary material for "Sub-clinical triiodothyronine levels predict health, demographic, and socioeconomic outcomes": Tables 1 and 2

| Table 1. Demographic, Socio-economic, and Health Relationships with Standardized Thyroid-Axis Hormones | | | | | | |  |  |  |  |  |  |  |  |  |
| --- | --- | --- | --- | --- | --- | --- | --- | --- | --- | --- | --- | --- | --- | --- | --- |
|  | [1] | [2] | [3] |  | [4] | [5] | [6] |  | [7] | [8] | [9] |  | [10] | [11] | [12] |
| Model Specification: | Base | | |  | Base & SES | | |  | Base, SES, Health and Health Behavior | | |  | Other Thyroid Hormones | | |
|  | Free T3 | Free T4 | TSH |  | Free T3 | Free T4 | TSH |  | Free T3 | Free T4 | TSH |  | Free T3 | Free T4 | TSH |
| Free T3 |  |  |  |  |  |  |  |  |  |  |  |  |  | 0.15*** | -0.02 |
|  |  |  |  |  |  |  |  |  |  |  |  |  |  | (0.02) | (0.01) |
| Free T4 |  |  |  |  |  |  |  |  |  |  |  |  | 0.14*** |  | -0.13*** |
|  |  |  |  |  |  |  |  |  |  |  |  |  | (0.02) |  | (0.01) |
| TSH |  |  |  |  |  |  |  |  |  |  |  |  | -0.02 | -0.12*** |  |
|  |  |  |  |  |  |  |  |  |  |  |  |  | (0.01) | (0.01) |  |
| (1) Male | 0.51*** | 0.07** | 0.02 |  | 0.51*** | 0.07** | 0.03 |  | 0.57*** | 0.04 | 0.04 |  | 0.57*** | -0.04 | 0.06 |
|  | (0.02) | (0.03) | (0.03) |  | (0.02) | (0.03) | (0.03) |  | (0.04) | (0.04) | (0.05) |  | (0.04) | (0.05) | (0.05) |
| (1) Black | -0.12*** | -0.04 | -0.39*** |  | -0.16*** | -0.04 | -0.40*** |  | -0.18*** | -0.07* | -0.39*** |  | -0.18*** | -0.09** | -0.40*** |
|  | (0.04) | (0.04) | (0.03) |  | (0.04) | (0.04) | (0.04) |  | (0.04) | (0.04) | (0.04) |  | (0.04) | (0.04) | (0.04) |
| (1) Hispanic | 0.19*** | 0.09** | -0.13*** |  | 0.12** | 0.03 | -0.09* |  | 0.07 | 0.06 | -0.10* |  | 0.06 | 0.03 | -0.09* |
|  | (0.03) | (0.04) | (0.03) |  | (0.05) | (0.05) | (0.05) |  | (0.05) | (0.05) | (0.05) |  | (0.05) | (0.05) | (0.05) |
| (1) Race non-white, Black, or Hispanic | -0.01 | 0.33*** | -0.16*** |  | -0.02 | 0.25*** | -0.12** |  | 0.04 | 0.22*** | -0.09 |  | 0.01 | 0.21*** | -0.06 |
|  | (0.06) | (0.06) | (0.04) |  | (0.07) | (0.05) | (0.05) |  | (0.08) | (0.05) | (0.06) |  | (0.08) | (0.05) | (0.06) |
| (1) Summer measurement | -0.09** | -0.02 | -0.00 |  | -0.10** | -0.02 | -0.00 |  | -0.11** | -0.01 | 0.01 |  | -0.11** | 0.00 | 0.00 |
|  | (0.04) | (0.06) | (0.03) |  | (0.04) | (0.06) | (0.03) |  | (0.05) | (0.06) | (0.03) |  | (0.05) | (0.06) | (0.03) |
| ln(Real HH Income) |  |  |  |  | -0.05*** | -0.01 | -0.03** |  | -0.04** | -0.01 | -0.01 |  | -0.04** | -0.01 | -0.01 |
|  |  |  |  |  | (0.02) | (0.02) | (0.01) |  | (0.02) | (0.02) | (0.01) |  | (0.02) | (0.02) | (0.01) |
| (1) College Ed. |  |  |  |  | -0.10*** | 0.07** | 0.02 |  | -0.04 | 0.06* | 0.04 |  | -0.05 | 0.07** | 0.05 |
|  |  |  |  |  | (0.04) | (0.03) | (0.04) |  | (0.04) | (0.03) | (0.05) |  | (0.04) | (0.03) | (0.05) |
| Constant | 0.31*** | -0.14* | -0.00 |  | 0.93*** | -0.07 | 0.35** |  | 0.84*** | -0.01 | 0.15 |  | 0.84*** | -0.12 | 0.16 |
|  | (0.06) | (0.07) | (0.04) |  | (0.19) | (0.21) | (0.16) |  | (0.20) | (0.21) | (0.17) |  | (0.19) | (0.21) | (0.17) |
| Observations | 8,059 | 8,059 | 8,059 |  | 8,059 | 8,059 | 8,059 |  | 7,426 | 7,426 | 7,426 |  | 7,426 | 7,426 | 7,426 |
| R-squared | 0.238 | 0.096 | 0.051 |  | 0.243 | 0.098 | 0.052 |  | 0.256 | 0.106 | 0.060 |  | 0.273 | 0.139 | 0.076 |
| Standard errors in parentheses. Models utilize weights to account for the population sampling probabilities of the NHANES, and use linearized standard errors. | | | | | | | | | | | |  |  |  |  |
| All models conditional on age, medication use, smoking, and wave. SES models conditional on household size and nativity. Health models conditional on Height, waist circumference, hours of sleep, iodine levels, calories per day, grams of sugar per day, and %HbA1c. T3, T4, and TSH outcomes expressed as standard deviations; *** p<0.01, ** p<0.05, * p<0.1 | | | | | | | | | | | | | | | |

| Table 2. Relationships between Free T3 and Labor Market Outcomes | | | | |  |  |  |  |  |  |  |  |  |  |  |  |  |
| --- | --- | --- | --- | --- | --- | --- | --- | --- | --- | --- | --- | --- | --- | --- | --- | --- | --- |
|  | [1] | [2] |  | [3] | [4] |  | [5] | [6] |  | [7] | [8] |  | [9] |  | [10] |  | [11] |
|  | Employed | | | | | | | | | | |  | Hours Worked | | | | |
|  | Base Model | |  | Base Model | |  | Add SES | |  | Add Health | |  | All Men |  | Employed Men |  | Employed in High-Activity Job |
|  | OLS | Logit |  | OLS | Logit |  | OLS | Logit |  | OLS | Logit |  | OLS |  | OLS |  | OLS |
| Free T3 | 0.01 | 1.03 |  | 0.02* | 1.05 |  | 0.01 | 1.06 |  | 0.01 | 1.04 |  | 0.13 |  | 0.56 |  | 1.53** |
|  | (0.01) | (0.06) |  | (0.01) | (0.07) |  | (0.01) | (0.07) |  | (0.01) | (0.07) |  | (0.57) |  | (0.59) |  | (0.72) |
| T3 * Age |  |  |  | 0.03*** | 1.18*** |  | 0.02*** | 1.12*** |  | 0.01** | 1.07* |  | 0.97*** |  | 1.10*** |  | 0.87** |
|  |  |  |  | (0.00) | (0.04) |  | (0.00) | (0.04) |  | (0.00) | (0.04) |  | (0.25) |  | (0.30) |  | (0.34) |
| Age (10 years) | -0.11*** | 0.55*** |  | -0.13*** | 0.46*** |  | -0.23** | 0.39 |  | -0.18* | 0.42 |  | -3.70 |  | 1.24 |  | -3.86 |
|  | (0.01) | (0.03) |  | (0.01) | (0.04) |  | (0.09) | (0.25) |  | (0.09) | (0.30) |  | (5.17) |  | (4.79) |  | (7.13) |
| Models utilize weights to account for the population sampling probabilities of the NHANES, and use linearized standard errors. | | | | | | | | | | |  |  |  |  |  |  |  |
| Coefficients come from a fully interacted model with age and the described covariates. Standard errors in parentheses. | | | | | | | | | | |  |  |  |  |  |  |  |
| T3, T4, and TSH outcomes expressed as standard deviations | | | | |  |  |  |  |  |  |  |  |  |  |  |  |  |
| *** p<0.01, ** p<0.05, * p<0.1 | |  |  |  |  |  |  |  |  |  |  |  |  |  |  |  |  |
