## Supplement B & Figures for "Sub-clinical triiodothyronine levels predict health, demographic, and socioeconomic outcomes"

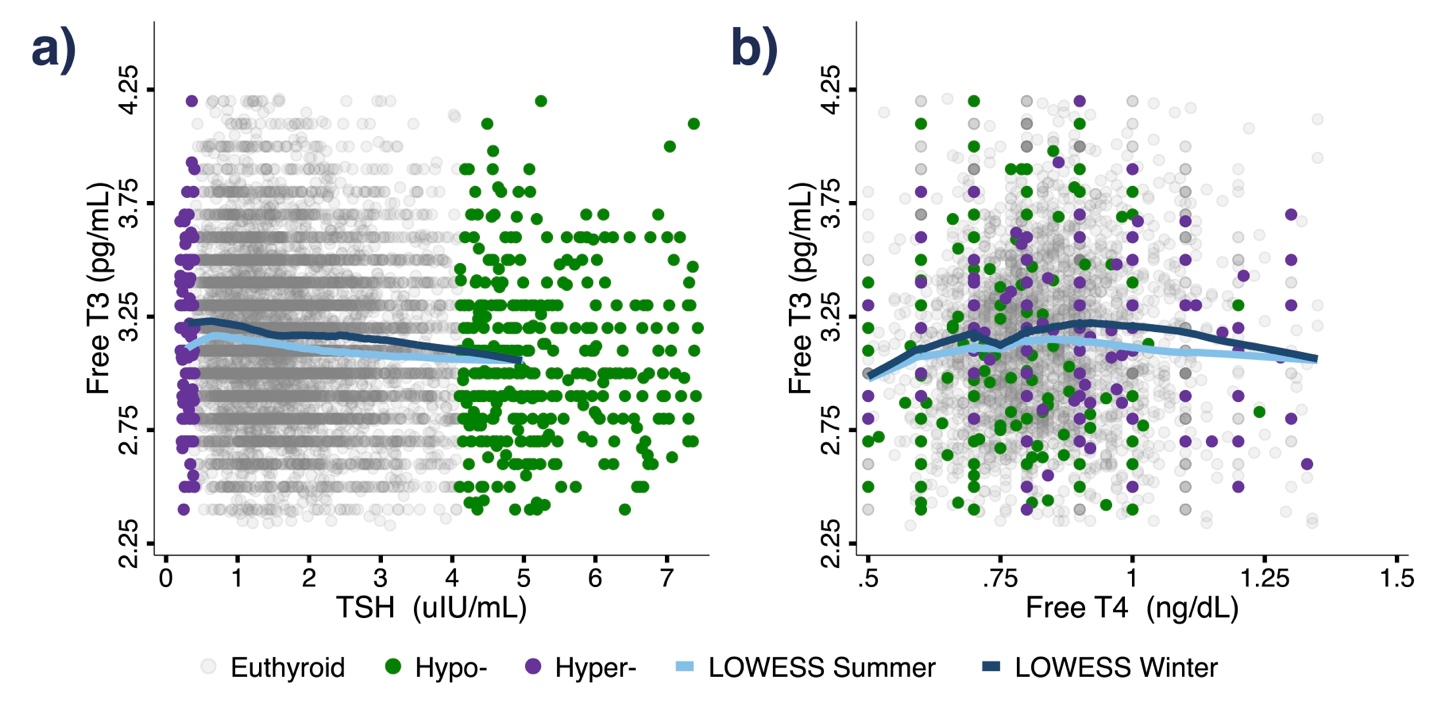

**Supplement Figure S1.** Inter-relationships between TSH, free T4, and free T3 in clinical units. a) Scatter plot showing TSH and free T3 for the same individual. b) Scatter plot showing free T4 and free T3 for the same individual. Euthyroid, hypo-, and hyper- thyroid individuals in both panels classified by TSH (hypo: TSH>4.1 mIU/L, hyper: TSH<0.4 mIU/L). Non-parametric LOWESS estimates shown stratifying by summer measurement (May 1-Oct. 31).

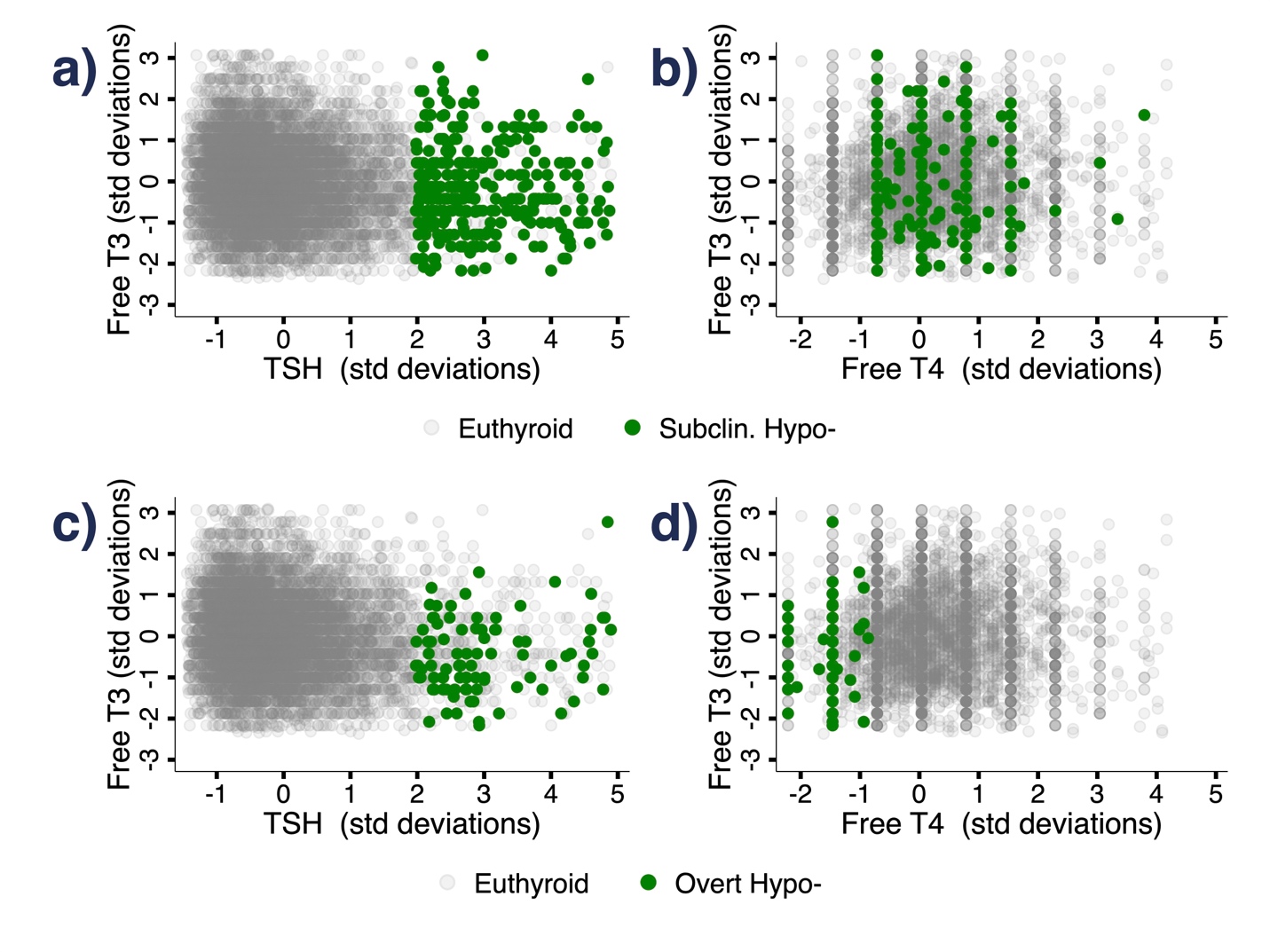

**Supplement Figure S2.** Inter-relationships between TSH, free T4, and free T3 in standard deviations, stratifying overt and subclinical hypothyroidism. a) Scatter plot showing TSH and free T3 for the same individual, in terms of standard deviations of the adult population distribution, highlighting subclinical hypothyroidism in green (TSH>4.1mIU/L, free T4>0.7ng/dL). b) Scatter plot showing free T4 and free T3 for the same individual, in terms of standard deviations of the adult population distribution, highlighting subclinical hypothyroidism in green. c) Scatter plot showing TSH and free T3 for the same individual, in terms of standard deviations of the adult population distribution, highlighting overt hypothyroidism in green (TSH>4.1mIU/L, free T4<0.7ng/dL). d) Scatter plot showing free T4 and free T3 for the same individual, in terms of standard deviations of the adult population distribution, highlighting overt hypothyroidism in green.

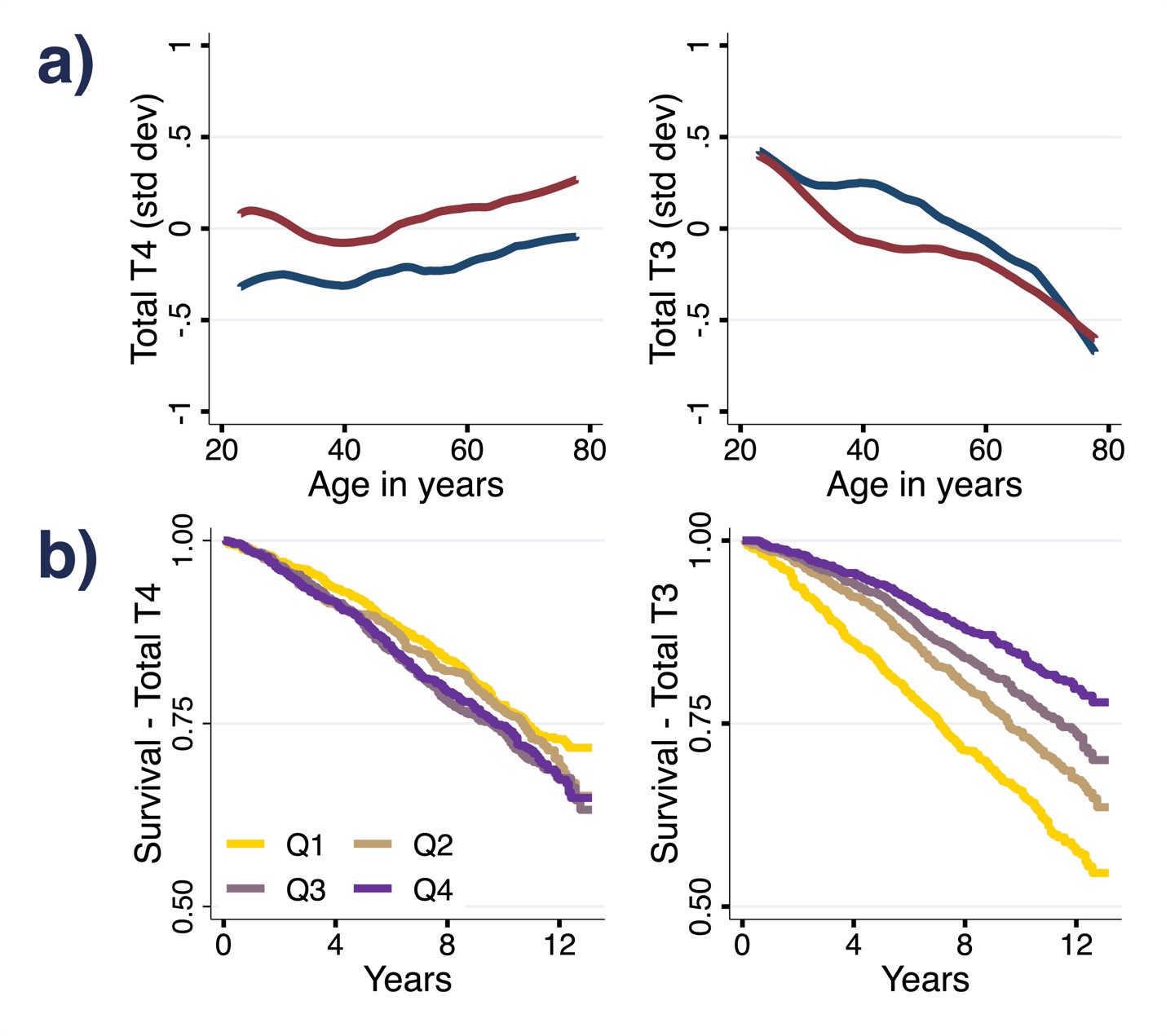

**Supplement Figure S3.** Total T4, Total T3, age, and mortality. **A)** weighted non-parametric LOWESS estimates of the relationships between thyroid hormones and age among adults, stratified by sex. **B)** weighted survival curves since measurement date for each quartile of total T4 and total T3, among adults over age 50 (Q1: 0-25%, Q2: 25-50%, Q3: 50-75%, Q4: 75-100% within each hormone’s distribution).

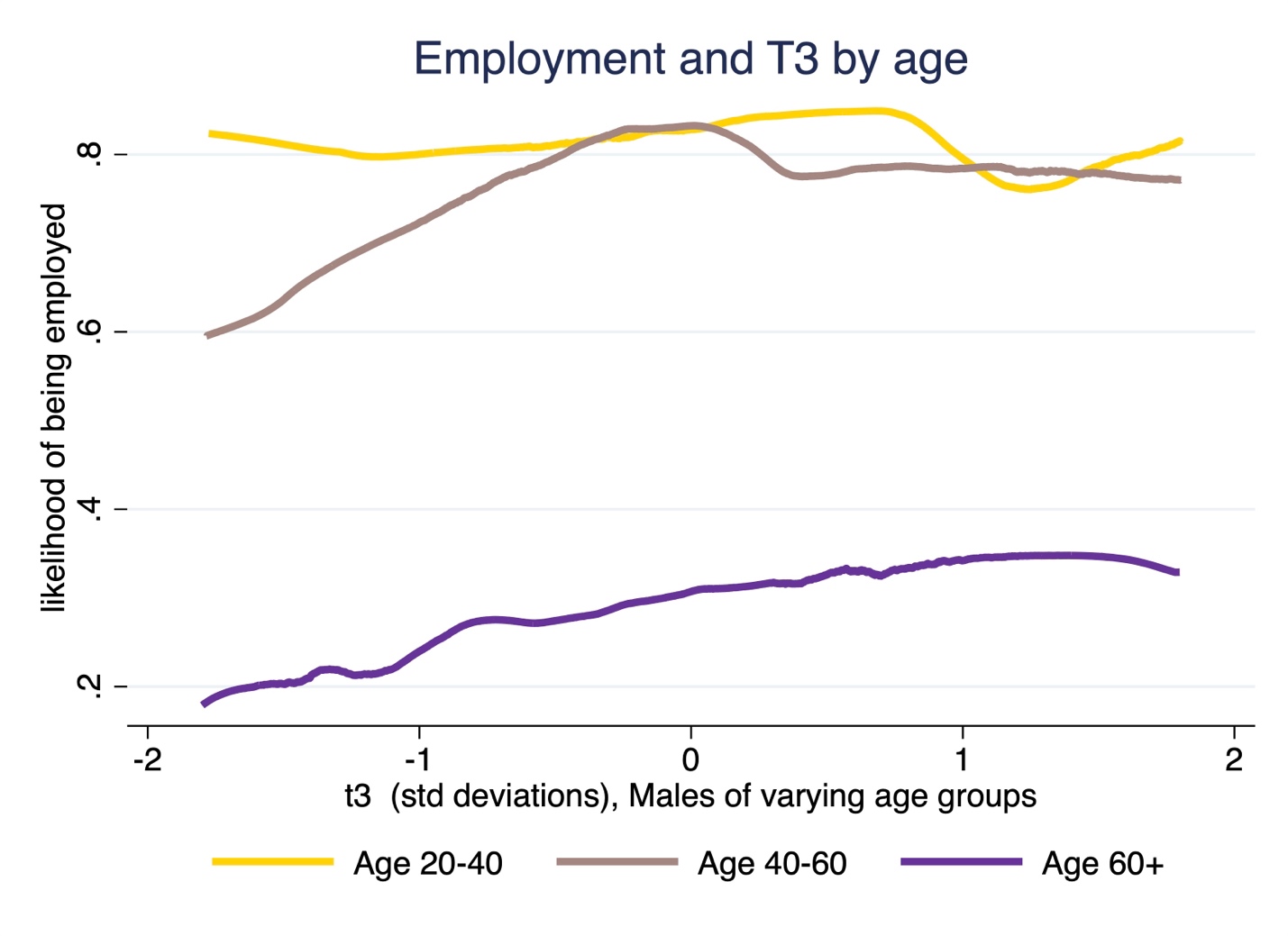

**Supplement Figure S4.** Employment likelihood and free T3 among males of varying age groups. Weighted non-parametric LOWESS estimates of the relationship between free T3 and employment status at the time of measurement are shown. T3 standard deviations are calculated within each age group set. Data from 2007-2012 NHANES.

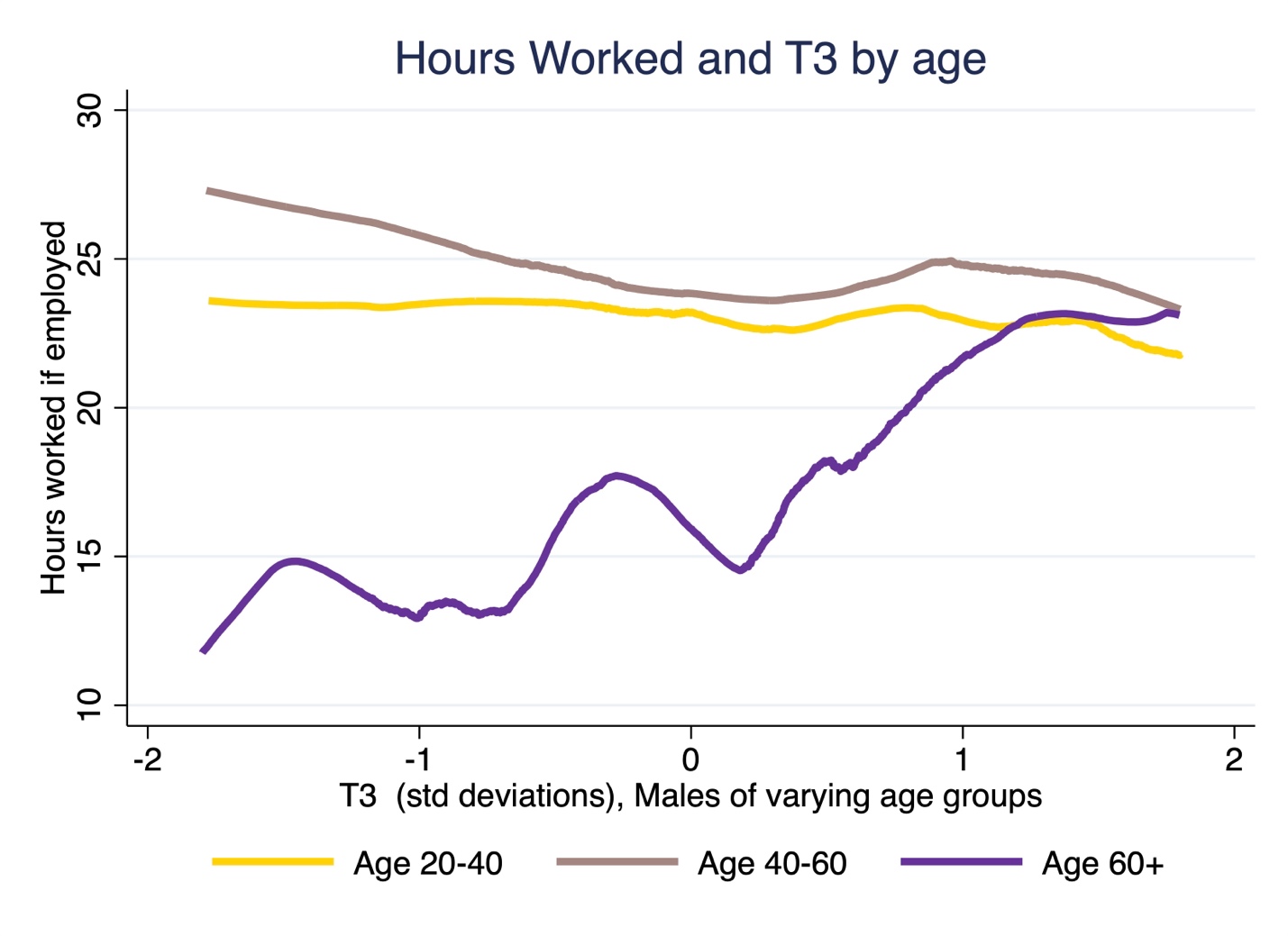

**Supplement Figure S5.** Hours worked and free T3 among males of varying age groups. Weighted non-parametric LOWESS estimates of the relationship between free T3 and hours worked among employed adults are shown. T3 standard deviations are calculated within each age group set. Data from 2007-2012 NHANES.

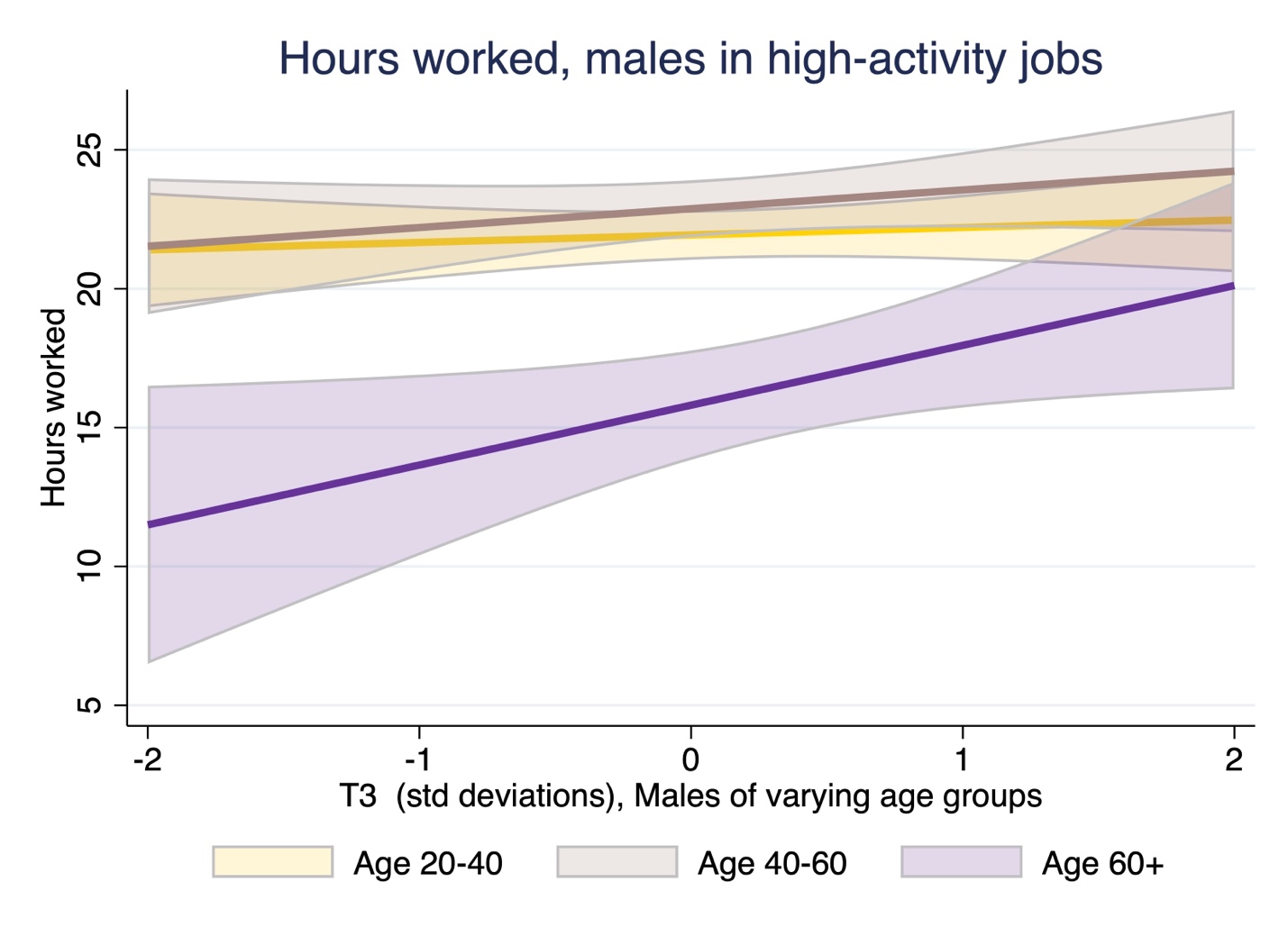

**Supplement Figure S6.** Hours worked and free T3 among males of varying age groups working in ‘high activity’ manual or service industry jobs. Weighted linear regression estimates of the relationship between free T3 and hours worked among employed adults are shown. 90% confidence intervals shown. Linear regression used instead of LOWESS due to restricted sample size. T3 standard deviations are calculated within each age group set. Data from 2007-2012 NHANES.

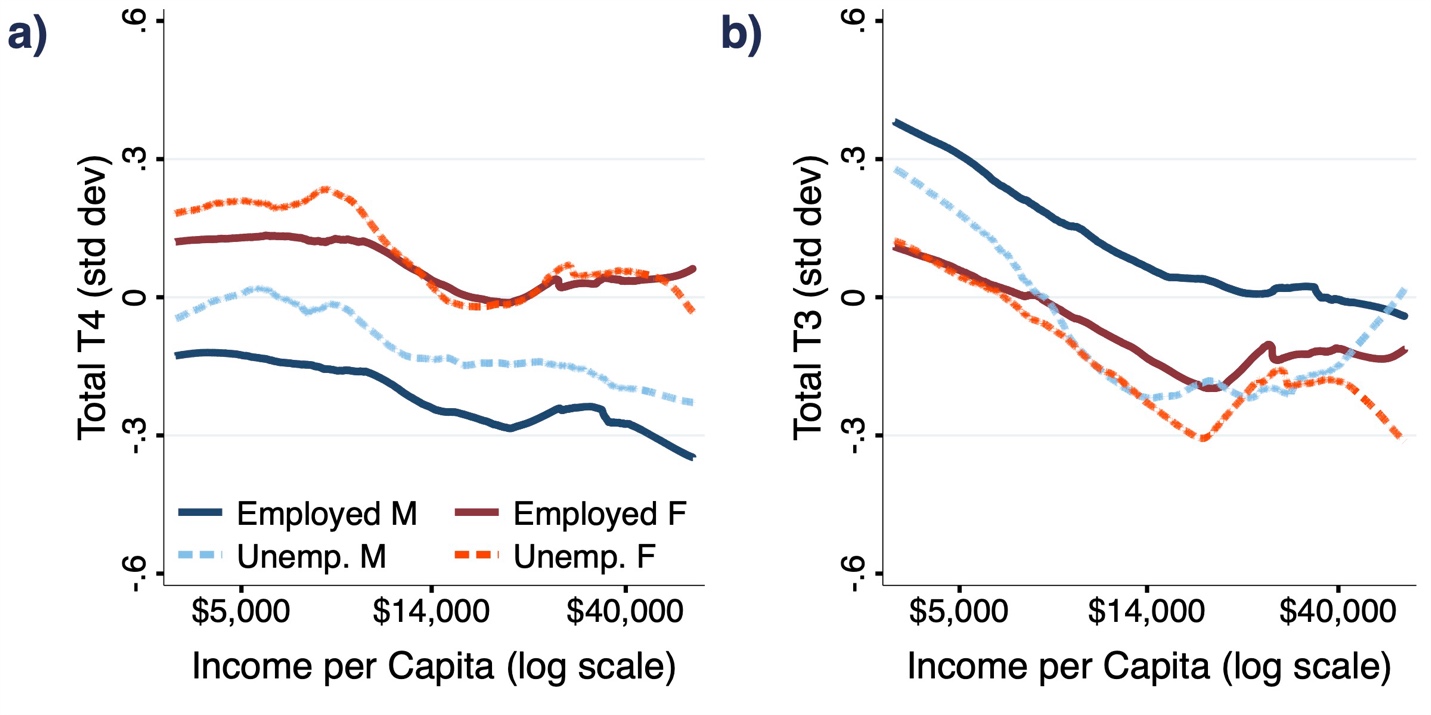

**Supplement Figure S7.** Total T4, Total T3, household income, and unemployment. Weighted non-parametric LOWESS estimates of the relationship between Total T4 and T3 and the natural logarithm of real (2007 dollars) household income per capita are displayed, stratified by sex and employment status at the time of measurement. Data from 2007-2012 NHANES.

Supplement B1. Regression-Kink Tests

Analysis for a regression kink were conducted following Hansen (2017). In the first-stage, visualized below, we minimize the least-squares criterion after iteratively estimating threshold regressions as specified below, where zi represents the free T3 outcome, yi represents the natural log of real income per capita, τ is the location of the income threshold (which we are trying to estimate), I(.) is an indicator function, β_1_ and β_2_ are slope parameters, and εi represents the error term.

(1)	zi =β0 +β_1_yi +β_2_(yi −τ)×I(yi −τ >0)+εi

We iteratively estimate (1) in increments of 0.01 with respect to the natural log of real income per capita, and plot the MSE in the figure below, identifying the point at which the least-squared error is minimized, in this case at $22,735.

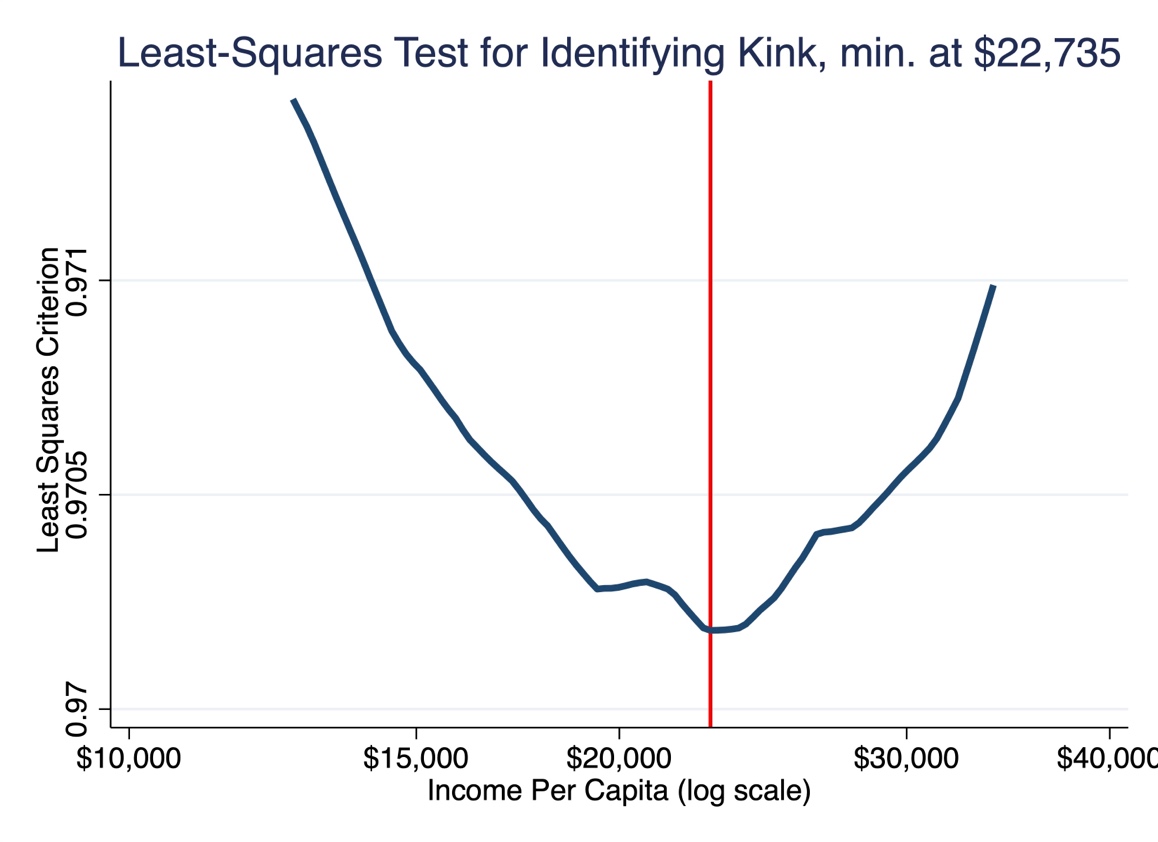

Having identified the optimal location for a kink, we first calculate an F-like statistic as described in Hansen (2017). In order to conduct inference on this statistic, we use a boot-strap approach with 1,000 replications that calculates simulated F-like statistics, and compare these to our calculated F-like statistic for our optimal threshold regression. In doing so, we find strong evidence rejecting the null hypothesis that β2 = 0, with p<0.001.
