## Supplemental Tables for "Sub-clinical triiodothyronine levels predict health, demographic, and socioeconomic outcomes"

| Supplement Table 1: Summary statistics from NHANES analytic sample, Age 20+ | | | | |  |  |  |  |  |
| --- | --- | --- | --- | --- | --- | --- | --- | --- | --- |
|  | Males | | | |  | Females | | | |
|  | Mean | SD | Max | Min |  | Mean | SD | Max | Min |
| TSH (uIU/mL) | 1.875 | (1.125) | 7.430 | 0.200 |  | 1.880 | (1.154) | 7.380 | 0.198 |
| Free T4 (ng/dL) | 0.791 | (0.127) | 0.500 | 0.796 |  | 0.792 | (0.134) | 1.350 | 0.500 |
| Free T3 (pg/mL) | 3.258 | (0.338) | 4.210 | 2.360 |  | 3.062 | (0.316) | 4.200 | 2.330 |
| Age in years | 46.87 | (15.93) | 80 | 21 |  | 48.25 | (16.40) | 80 | 21 |
| ln(real householdincome) | 10.75 | (0.754) | 11.65 | 7.743 |  | 10.64 | (0.817) | 11.65 | 7.743 |
| Proportion on thyroid medication | 0.0208 | (0.143) | - | - |  | 0.0934 | (0.291) | - | - |
| Proportion Employed | 0.683 | (0.465) | - | - |  | 0.548 | (0.498) | - | - |
| Proportion Black | 0.0932 | (0.291) | - | - |  | 0.111 | (0.315) | - | - |
| Proportion Hispanic | 0.143 | (0.350) | - | - |  | 0.128 | (0.334) | - | - |
| Proportion measured in summer | 0.601 | (0.490) | - | - |  | 0.617 | (0.486) | - | - |
| Height (m) | 1.759 | (0.0763) | 2.045 | 1.421 |  | 1.622 | (0.0703) | 1.942 | 1.345 |
| Waist Circumference (m) | 1.014 | (0.149) | 1.782 | 0.654 |  | 0.953 | (0.157) | 1.703 | 0.591 |
| Proportion wave 2009-2010 | 0.205 | (0.404) | - | - |  | 0.200 | (0.400) | - | - |
| Proportion wave 2011-2012 | 0.204 | (0.403) | - | - |  | 0.208 | (0.406) | - | - |
| Proportion high school completed | 0.261 | (0.439) | - | - |  | 0.258 | (0.438) | - | - |
| Proportion some college | 0.281 | (0.449) | - | - |  | 0.279 | (0.449) | - | - |
| Proportion college completed | 0.278 | (0.448) | - | - |  | 0.274 | (0.446) | - | - |
| Sample Size | 4,087 | | | |  | 3,972 | | | |
| Models utilize weights to account for the population sampling probabilities of the NHANES. | | | | | |  |  |  |  |

| Supplement Table S2. Relationships Between HPT-axis Hormones and Clinical Cutoffs | | | | | | |  |  |  |  |  |  |  |
| --- | --- | --- | --- | --- | --- | --- | --- | --- | --- | --- | --- | --- | --- |
|  | [1] | [2] | [3] |  | [4] | [5] | [6] |  | [7] | [8] |  | [9] | [10] |
|  | Free T3 | | | | | | | | | | | | |
|  | All Adults | | | | | | |  | All Adults | |  | Adults not on Meds | |
|  | No Controls | | |  | Base Model | | |  | No Controls | Base Model |  | No Controls | Base Model |
| Free T4 | 0.06*** |  | 0.05*** |  | 0.13*** |  | 0.12*** |  |  |  |  |  |  |
|  | (0.01) |  | (0.01) |  | (0.01) |  | (0.01) |  |  |  |  |  |  |
| TSH |  | -0.09*** | -0.08*** |  |  | -0.03** | -0.01 |  |  |  |  |  |  |
|  |  | (0.01) | (0.01) |  |  | (0.01) | (0.01) |  |  |  |  |  |  |
| Hypothyroid |  |  |  |  |  |  |  |  | -0.53*** | -0.35*** |  | -0.51*** | -0.31*** |
|  |  |  |  |  |  |  |  |  | (0.11) | (0.09) |  | (0.11) | (0.10) |
| Subclinical Hypothyroid |  |  |  |  |  |  |  |  | -0.17* | -0.00 |  | -0.08 | 0.01 |
|  |  |  |  |  |  |  |  |  | (0.09) | (0.07) |  | (0.09) | (0.08) |
| Hyperthyroid |  |  |  |  |  |  |  |  | -0.20* | -0.02 |  | -0.03 | -0.01 |
|  |  |  |  |  |  |  |  |  | (0.10) | (0.09) |  | (0.10) | (0.10) |
| R-squared | 0.004 | 0.008 | 0.010 |  | 0.252 | 0.239 | 0.252 |  | 0.005 | 0.240 |  | 0.003 | 0.217 |
| Models utilize weights to account for the population sampling probabilities of the NHANES, and use linearized standard errors. | | | | | | | | | | |  |  |  |
| Adults not on meds excludes all adults on thyroid-related medications. | | | | | |  |  |  |  |  |  |  |  |
| Clinical category definitions: Hypothyroid (TSH>4.1 & Free T4<15th %ile), Subclinical Hypothyroid (TSH>4.1 & T4>15th %ile), Hyperthyroid (TSH<0.4) | | | | | | | | | | | | |  |
| T3, T4, and TSH outcomes expressed as standard deviations | | | |  |  |  |  |  |  |  |  |  |  |
| *** p<0.01, ** p<0.05, * p<0.1 | |  |  |  |  |  |  |  |  |  |  |  |  |

| Supplement Table S3. Relationships between Thyroid Hormones and Mortality, Age 50+ | | | | | | | | |
| --- | --- | --- | --- | --- | --- | --- | --- | --- |
|  | [1] | [2] | [3] | [4] |  | [5] |  | [6] |
|  | Base Model | | | |  | Add SES |  | Add Health |
|  | Mortality | | | | | | | |
| Free T3 | 0.87*** |  |  | 0.86*** |  | 0.85*** |  | 0.89** |
|  | (0.04) |  |  | (0.04) |  | (0.04) |  | (0.05) |
| Free T4 |  | 1.19*** |  | 1.21*** |  | 1.22*** |  | 1.21*** |
|  |  | (0.04) |  | (0.04) |  | (0.04) |  | (0.05) |
| TSH |  |  | 1.02 | 1.05 |  | 1.05 |  | 1.05 |
|  |  |  | (0.03) | (0.03) |  | (0.03) |  | (0.04) |
| Models utilize weights to account for the population sampling probabilities of the NHANES, and use linearized standard errors. | | | | | | | | |
| Hazard Ratios come from a Cox proportional hazards model conditional on the described covariates. Standard errors in parentheses. | | | | | | | | |
| Sample restricted to over 50 years old to focus on mortality. Mortality data through 2019. | | | | | | | | |
| T3, T4, and TSH outcomes expressed as standard deviations | | | | |  |  |  |  |
| *** p<0.01, ** p<0.05, * p<0.1 | | |  |  |  |  |  |  |

| Supplement Table S4. Relationships between Total T3, Total T4, and Mortality, Age 50+ | | | | | | |
| --- | --- | --- | --- | --- | --- | --- |
|  | [1] | [2] |  | [5] |  | [6] |
|  | Base Model | |  | Add SES + Health | | |
|  | Mortality | | | | | |
| Total T3 | 0.84*** |  |  | 0.87*** |  |  |
|  | (0.04) |  |  | (0.04) |  |  |
| Total T4 |  | 1.14*** |  |  |  | 1.09** |
|  |  | (0.05) |  |  |  | (0.05) |
| Models utilize weights to account for the population sampling probabilities of the NHANES, and use linearized standard errors. | | | | | | |
| Hazard Ratios come from a Cox proportional hazards model conditional on the described covariates. Standard errors in parentheses. | | | | | | |
| Sample restricted to over 50 years old to focus on mortality. Mortality data through 2019. | | | | | | |
| Total T3 and T4 expressed as standard deviations | | | | |  |  |
| *** p<0.01, ** p<0.05, * p<0.1 | | |  |  |  |  |

| Supplement Table S5. Relationships between Thyroid Hormones, Mortality, and Illness, Age 50+ | | |
| --- | --- | --- |
|  | [1] | [2] |
|  | Full Model with Illness Covariates | |
|  | w/o CRP | with CRP |
|  | Mortality | |
| Free T3 | 0.89** | 0.91* |
|  | (0.05) | (0.05) |
| Free T4 | 1.21*** | 1.18*** |
|  | (0.05) | (0.05) |
| TSH | 1.05 | 1.02 |
|  | (0.04) | (0.05) |
| Observations | 3,633 | 3,019 |
| Models utilize weights to account for the population sampling probabilities of the NHANES, and use linearized standard errors. | | |
| Hazard Ratios come from a Cox proportional hazards model conditional on the described covariates. Standard errors in parentheses. | | |
| C-Reactive Protein only in first two waves, reducing sample size | | |
| Ilnness covariates include whether or not respondent has had a cold, gastrointestinal illness, or flu within the past 30 days. | | |
| Sample restricted to over 50 years old to focus on mortality. Mortality data through 2019. | | |
| T3, T4, and TSH outcomes expressed as standard deviations | | |
| *** p<0.01, ** p<0.05, * p<0.1 | | |

| Supplement Table S6. Alternate Income Specifications for Demographic, Socio-economic, and Health Relationships with Standardized Thyroid-Axis Hormones | | | | | | | | |
| --- | --- | --- | --- | --- | --- | --- | --- | --- |
|  | [1] | [2] |  | [3] | [4] |  | [5] | [6] |
| Model Specification: | Per-Capita Income | Income-Poverty Ratio |  | Per-Capita Income | Income-Poverty Ratio |  | Per-Capita Income | Income-Poverty Ratio |
|  | Free T3 | Free T3 |  | Free T4 | Free T4 |  | TSH | TSH |
| ln(Real Per Capita HH Income) | -0.04** |  |  | -0.02 |  |  | -0.01 |  |
|  | (0.02) |  |  | (0.02) |  |  | (0.01) |  |
| Poverty Index Ratio |  | -0.02* |  |  | -0.00 |  |  | -0.01 |
|  |  | (0.01) |  |  | (0.01) |  |  | (0.01) |
| (1) Male | 0.57*** | 0.57*** |  | 0.04 | 0.05 |  | 0.04 | 0.04 |
|  | (0.04) | (0.04) |  | (0.04) | (0.05) |  | (0.05) | (0.05) |
| (1) Age 31-40 | -0.29*** | -0.30*** |  | -0.17*** | -0.17*** |  | 0.07 | 0.05 |
|  | (0.04) | (0.05) |  | (0.05) | (0.05) |  | (0.04) | (0.04) |
| (1) Age 41-50 | -0.47*** | -0.46*** |  | -0.25*** | -0.25*** |  | 0.16*** | 0.16*** |
|  | (0.05) | (0.06) |  | (0.05) | (0.05) |  | (0.05) | (0.06) |
| (1) Age 51-60 | -0.57*** | -0.58*** |  | -0.17** | -0.15** |  | 0.21*** | 0.20*** |
|  | (0.04) | (0.05) |  | (0.07) | (0.07) |  | (0.05) | (0.06) |
| (1) Age 61-70 | -0.82*** | -0.84*** |  | 0.05 | 0.06 |  | 0.19*** | 0.17*** |
|  | (0.06) | (0.06) |  | (0.06) | (0.06) |  | (0.06) | (0.06) |
| (1) Age 71+ | -1.15*** | -1.17*** |  | 0.18** | 0.17** |  | 0.41*** | 0.39*** |
|  | (0.06) | (0.06) |  | (0.08) | (0.08) |  | (0.06) | (0.07) |
| (1) Black | -0.18*** | -0.17*** |  | -0.07* | -0.06 |  | -0.39*** | -0.39*** |
|  | (0.04) | (0.04) |  | (0.04) | (0.04) |  | (0.04) | (0.04) |
| (1) Hispanic | 0.08 | 0.07 |  | 0.05 | 0.06 |  | -0.10* | -0.11** |
|  | (0.05) | (0.05) |  | (0.05) | (0.06) |  | (0.05) | (0.05) |
| (1) Race non-white, Black, or Hispanic | 0.04 | 0.03 |  | 0.22*** | 0.21*** |  | -0.09 | -0.07 |
|  | (0.08) | (0.08) |  | (0.05) | (0.06) |  | (0.06) | (0.07) |
| (1) Takes any thyroid medication | -0.35*** | -0.35*** |  | 0.87*** | 0.85*** |  | 0.12 | 0.16 |
|  | (0.05) | (0.06) |  | (0.08) | (0.08) |  | (0.09) | (0.10) |
| (1) Takes Statin | -0.07** | -0.08** |  | -0.02 | -0.02 |  | -0.07 | -0.10* |
|  | (0.03) | (0.03) |  | (0.04) | (0.04) |  | (0.04) | (0.05) |
| (1) Summer measurement | -0.11** | -0.12** |  | -0.01 | -0.02 |  | 0.01 | -0.01 |
|  | (0.05) | (0.04) |  | (0.06) | (0.06) |  | (0.03) | (0.03) |
| (1) College Ed. | -0.05 | -0.05 |  | 0.06* | 0.06* |  | 0.04 | 0.06 |
|  | (0.04) | (0.04) |  | (0.03) | (0.03) |  | (0.05) | (0.05) |
| Constant | 0.39*** | 0.44*** |  | -0.15* | -0.13 |  | 0.02 | 0.05 |
|  | (0.08) | (0.08) |  | (0.08) | (0.08) |  | (0.05) | (0.06) |
| Observations | 7,426 | 6,799 |  | 7,426 | 6,799 |  | 7,426 | 6,799 |
| R-squared | 0.256 | 0.257 |  | 0.106 | 0.105 |  | 0.060 | 0.060 |
| Standard errors in parentheses. Models utilize weights to account for the population sampling probabilities of the NHANES, and use linearized standard errors. | | | | | | | | |
| Poverty Index Ratio is calculated using HHS guidelines, dividing household income by household size, state, and year specific poverty thresholds | | | | | | | | |
| All models conditional on age, medication use, smoking, and wave. SES Models consistent on household size and nativity. Health models conditional on Height, waist circumference, hours of sleep, iodine levels, calories per day, grams of sugar per day, and %HbA1c. | | | | | | | | |

| Supplement Table S7. SES Regressions of Other Biomarkers (Standardized) | | | |  |  |  |
| --- | --- | --- | --- | --- | --- | --- |
|  | [1] | [2] | [3] | [4] | [5] | [6] |
| Model Specification: | Base & SES | | | | | |
|  | HDL | Non-HDL | %HbA1C | Sys. BP | Dias. BP | CRP |
| (1) Male | -0.41*** | 0.11*** | 0.03*** | 0.25*** | 0.26*** | -0.17*** |
|  | (0.02) | (0.02) | (0.01) | (0.02) | (0.02) | (0.03) |
| (1) Black | 0.15*** | -0.18*** | 0.14*** | 0.19*** | 0.07* | 0.19*** |
|  | (0.02) | (0.03) | (0.02) | (0.04) | (0.04) | (0.05) |
| (1) Hispanic | -0.07** | 0.03 | 0.10*** | 0.04 | -0.06 | 0.10* |
|  | (0.03) | (0.03) | (0.02) | (0.03) | (0.04) | (0.05) |
| (1) Race non-white, Black, or Hispanic | -0.07* | -0.06 | 0.06*** | 0.00 | 0.02 | -0.13*** |
|  | (0.03) | (0.05) | (0.02) | (0.04) | (0.06) | (0.04) |
| (1) Summer measurement | -0.04* | -0.00 | 0.01 | 0.03 | 0.03 | -0.02 |
|  | (0.02) | (0.02) | (0.01) | (0.02) | (0.03) | (0.03) |
| ln(Real HH Income) | 0.05*** | -0.02 | -0.04*** | -0.03 | 0.03 | -0.08*** |
|  | (0.01) | (0.01) | (0.01) | (0.02) | (0.02) | (0.03) |
| (1) College Ed. | 0.14*** | -0.10*** | -0.06*** | -0.16*** | -0.07** | -0.10*** |
|  | (0.02) | (0.02) | (0.01) | (0.03) | (0.03) | (0.03) |
| Constant | -0.76*** | -0.43*** | -0.34*** | -0.25 | -0.55** | 0.74*** |
|  | (0.13) | (0.16) | (0.08) | (0.18) | (0.25) | (0.27) |
| Observations | 8,058 | 8,058 | 8,045 | 7,501 | 7,501 | 6,563 |
| R-squared | 0.153 | 0.120 | 0.158 | 0.224 | 0.125 | 0.033 |
| Standard errors in parentheses. Models utilize weights to account for the population sampling probabilities of the NHANES, and use linearized standard errors. | | | | | | |
| All models conditional on age, medication use, smoking, household size, nativity, and wave. | | | | | |  |
| All biomarkers standardized so outcomes represent changes in number of standard deviations of a given biomarker | | | | | | |

| Supplement Table S8. Demographic, Socio-economic, and Health Relationships with Standardized Total T3 | | | | | |  |
| --- | --- | --- | --- | --- | --- | --- |
|  | [1] | [2] | [3] | [4] | [5] | [6] |
| Model Specification: | Total T3 | | | Total T4 | | |
|  | Base | SES | Full | Base | SES | Full |
| (1) Male | 0.11*** | 0.11*** | 0.19*** | -0.25*** | -0.24*** | -0.18*** |
|  | (0.02) | (0.03) | (0.04) | (0.02) | (0.02) | (0.04) |
| (1) Black | -0.10** | -0.14*** | -0.16*** | 0.10** | 0.08 | 0.01 |
|  | (0.05) | (0.05) | (0.05) | (0.05) | (0.05) | (0.05) |
| (1) Hispanic | 0.16*** | 0.09** | 0.05 | 0.25*** | 0.16*** | 0.11** |
|  | (0.04) | (0.04) | (0.04) | (0.04) | (0.05) | (0.05) |
| (1) Race non-white, Black, or Hispanic | -0.14** | -0.14* | -0.11 | 0.21*** | 0.15*** | 0.14** |
|  | (0.06) | (0.07) | (0.08) | (0.04) | (0.05) | (0.05) |
| (1) Summer measurement | -0.03 | -0.04 | -0.06 | -0.17*** | -0.17*** | -0.18*** |
|  | (0.05) | (0.05) | (0.05) | (0.05) | (0.05) | (0.05) |
| ln(Real HH Income) |  | -0.05** | -0.04* |  | -0.04** | -0.02 |
|  |  | (0.02) | (0.02) |  | (0.02) | (0.02) |
| (1) College Ed. |  | -0.14*** | -0.10** |  | -0.04 | -0.00 |
|  |  | (0.04) | (0.04) |  | (0.03) | (0.03) |
| Constant | 0.24*** | 0.79*** | 0.71*** | 0.00 | 0.46** | 0.35* |
|  | (0.07) | (0.25) | (0.24) | (0.05) | (0.19) | (0.18) |
| Observations | 8,059 | 8,059 | 7,426 | 8,031 | 8,031 | 7,399 |
| R-squared | 0.100 | 0.106 | 0.126 | 0.071 | 0.073 | 0.099 |
| Standard errors in parentheses. Models utilize weights to account for the population sampling probabilities of the NHANES, and use linearized standard errors. | | | | | | |
| All models conditional on age, medication use, smoking, and wave. SES models conditional on household size and nativity. Health models conditional on Height, waist circumference, hours of sleep, iodine levels, calories per day, grams of sugar per day, and %HbA1c. | | | | | | |

| Supplement Table S9. Relationships between Total T3 and Labor Market Outcomes | | | | | | | |  |  |  |  |  |  |  |  |  |  |
| --- | --- | --- | --- | --- | --- | --- | --- | --- | --- | --- | --- | --- | --- | --- | --- | --- | --- |
|  | [1] | [2] |  | [3] | [4] |  | [5] | [6] |  | [7] | [8] |  | [9] |  | [10] |  | [11] |
|  | Employed | | | | | | | | | | |  | Hours Worked | | | | |
|  | Base Model | |  | Base Model | |  | Add SES | |  | Add Health | |  | All Men |  | Employed Men |  | Employed in High-Activity Job |
|  | OLS | Logit |  | OLS | Logit |  | OLS | Logit |  | OLS | Logit |  | OLS |  | OLS |  | OLS |
| Free T3 | 0.01 | 1.02 |  | 0.01 | 1.01 |  | 0.01 | 1.03 |  | 0.00 | 1.01 |  | 0.03 |  | 0.29 |  | 0.39 |
|  | (0.01) | (0.07) |  | (0.01) | (0.07) |  | (0.01) | (0.06) |  | (0.01) | (0.06) |  | (0.55) |  | (0.52) |  | (0.73) |
| T3 * Age |  |  |  | 0.02*** | 1.13*** |  | 0.01** | 1.09* |  | 0.01 | 1.06 |  | 0.51 |  | 0.56 |  | 0.02 |
|  |  |  |  | (0.01) | (0.04) |  | (0.01) | (0.05) |  | (0.01) | (0.05) |  | (0.33) |  | (0.34) |  | (0.59) |
| Age (10 years) |  |  |  | -0.12*** | 0.48*** |  | 0.01** | 1.09* |  | -0.18* | 0.44 |  | -3.50 |  | 2.06 |  | -3.17 |
|  |  |  |  | (0.01) | (0.04) |  | (0.01) | (0.05) |  | (0.10) | (0.31) |  | (5.25) |  | (4.81) |  | (6.84) |
| Models utilize weights to account for the population sampling probabilities of the NHANES, and use linearized standard errors. | | | | | | | | | | | |  |  |  |  |  |  |
| Coefficients come from a fully interacted model with age and the described covariates. Standard errors in parentheses. | | | | | | | | | | |  |  |  |  |  |  |  |
| T3, T4, and TSH outcomes expressed as standard deviations | | | | |  |  |  |  |  |  |  |  |  |  |  |  |  |
| *** p<0.01, ** p<0.05, * p<0.1 | |  |  |  |  |  |  |  |  |  |  |  |  |  |  |  |  |

| Supplement S10. Robustness of Demographic, Socio-economic, and Health Relationships to Winsorization | | | | | |  |  |  |  |  |  |  |  |  |  |  |  |  |  |  |
| --- | --- | --- | --- | --- | --- | --- | --- | --- | --- | --- | --- | --- | --- | --- | --- | --- | --- | --- | --- | --- |
|  | [1] | [2] | [3] | [4] | [5] | [6] |  | [7] | [8] | [9] | [10] | [11] | [12] |  | [13] | [14] | [15] | [16] | [17] | [18] |
| Model Specification: | Free T3 | | | | | |  | Free T4 | | | | | |  | TSH | | | | | |
|  | Base - 6% | Base - 10% | SES - 6% | SES - 10% | Health - 6% | Health - 10% |  | Base - 6% | Base - 10% | SES - 6% | SES - 10% | Health - 6% | Health - 10% |  | Base - 6% | Base - 10% | SES - 6% | SES - 10% | Health - 6% | Health - 10% |
| (1) Male | 0.44*** | 0.42*** | 0.45*** | 0.42*** | 0.49*** | 0.46*** |  | 0.08*** | 0.07*** | 0.07*** | 0.07*** | 0.06* | 0.06* |  | 0.00 | -0.00 | 0.01 | 0.00 | 0.01 | 0.01 |
|  | (0.02) | (0.02) | (0.02) | (0.02) | (0.04) | (0.04) |  | (0.02) | (0.02) | (0.02) | (0.02) | (0.04) | (0.03) |  | (0.02) | (0.02) | (0.02) | (0.02) | (0.04) | (0.04) |
| (1) Black | -0.08*** | -0.08*** | -0.12*** | -0.11*** | -0.14*** | -0.13*** |  | -0.02 | -0.03 | -0.01 | -0.02 | -0.03 | -0.04 |  | -0.30*** | -0.27*** | -0.31*** | -0.27*** | -0.31*** | -0.28*** |
|  | (0.03) | (0.03) | (0.03) | (0.03) | (0.03) | (0.03) |  | (0.03) | (0.03) | (0.04) | (0.03) | (0.04) | (0.03) |  | (0.02) | (0.02) | (0.02) | (0.02) | (0.02) | (0.02) |
| (1) Hispanic | 0.18*** | 0.18*** | 0.13*** | 0.14*** | 0.10** | 0.11*** |  | 0.11*** | 0.08** | 0.06 | 0.04 | 0.08 | 0.05 |  | -0.10*** | -0.10*** | -0.06* | -0.07* | -0.07* | -0.09** |
|  | (0.03) | (0.03) | (0.04) | (0.04) | (0.04) | (0.04) |  | (0.04) | (0.04) | (0.05) | (0.04) | (0.05) | (0.05) |  | (0.02) | (0.02) | (0.03) | (0.03) | (0.04) | (0.04) |
| (1) Race non-white, Black, or Hispanic | 0.01 | 0.02 | 0.01 | 0.02 | 0.05 | 0.07 |  | 0.30*** | 0.27*** | 0.23*** | 0.20*** | 0.22*** | 0.19*** |  | -0.14*** | -0.13*** | -0.09** | -0.08 | -0.07 | -0.06 |
|  | (0.06) | (0.05) | (0.07) | (0.06) | (0.07) | (0.06) |  | (0.05) | (0.04) | (0.04) | (0.04) | (0.04) | (0.04) |  | (0.04) | (0.04) | (0.05) | (0.05) | (0.05) | (0.05) |
| (1) Summer measurement | -0.06 | -0.06 | -0.07* | -0.07* | -0.08* | -0.08* |  | -0.01 | -0.02 | -0.00 | -0.01 | -0.00 | -0.01 |  | 0.00 | -0.01 | 0.00 | -0.01 | 0.01 | 0.00 |
|  | (0.04) | (0.04) | (0.04) | (0.04) | (0.04) | (0.04) |  | (0.05) | (0.05) | (0.05) | (0.05) | (0.05) | (0.05) |  | (0.02) | (0.03) | (0.02) | (0.03) | (0.02) | (0.03) |
| ln(Real HH Income) |  |  | -0.05*** | -0.05*** | -0.04*** | -0.04*** |  |  |  | -0.00 | 0.00 | -0.01 | -0.00 |  |  |  | -0.01 | -0.01 | -0.00 | 0.00 |
|  |  |  | (0.01) | (0.01) | (0.01) | (0.01) |  |  |  | (0.02) | (0.02) | (0.02) | (0.02) |  |  |  | (0.01) | (0.01) | (0.01) | (0.01) |
| (1) College Ed. |  |  | -0.07** | -0.07** | -0.02 | -0.02 |  |  |  | 0.06** | 0.07** | 0.05 | 0.06** |  |  |  | -0.01 | -0.02 | -0.00 | -0.01 |
|  |  |  | (0.03) | (0.03) | (0.03) | (0.03) |  |  |  | (0.03) | (0.03) | (0.03) | (0.03) |  |  |  | (0.03) | (0.03) | (0.04) | (0.03) |
| Constant | 0.19*** | 0.21*** | 0.75*** | 0.74*** | 0.66*** | 0.64*** |  | -0.19*** | -0.21*** | -0.18 | -0.25 | -0.12 | -0.21 |  | -0.09*** | -0.09*** | 0.06 | 0.01 | -0.04 | -0.07 |
|  | (0.05) | (0.06) | (0.15) | (0.15) | (0.15) | (0.16) |  | (0.06) | (0.06) | (0.20) | (0.21) | (0.21) | (0.22) |  | (0.03) | (0.03) | (0.13) | (0.13) | (0.15) | (0.16) |
| Observations | 7,714 | 7,517 | 7,714 | 7,517 | 7,100 | 6,924 |  | 7,764 | 7,585 | 7,764 | 7,585 | 7,161 | 6,995 |  | 7,660 | 7,417 | 7,660 | 7,417 | 7,057 | 6,826 |
| R-squared | 0.219 | 0.199 | 0.223 | 0.203 | 0.233 | 0.215 |  | 0.075 | 0.068 | 0.078 | 0.071 | 0.086 | 0.080 |  | 0.049 | 0.048 | 0.050 | 0.049 | 0.055 | 0.054 |
| Standard errors in parentheses. Models utilize weights to account for the population sampling probabilities of the NHANES, and use linearized standard errors. | | | | | | | | | | |  |  |  |  |  |  |  |  |  |  |
| Robustness of estimates to alternative samples with different levels of winsorization tested, 6% (top and bottom 3%) and 10% (top and bottom 10%). | | | | | | | | | |  |  |  |  |  |  |  |  |  |  |  |
| All models conditional on age, medication use, smoking, and wave. SES models conditional on household size and nativity. Health models conditional on Height, waist circumference, hours of sleep, iodine levels, calories per day, grams of sugar per day, and %HbA1c. | | | | | | | | | | | | | | | | | | | |  |

| Supplement S11. Robustness of Relationships between Free T3 and likelihood of being unemployed to winsorization | | | | | | | | | | |  |  |  |  |  |  |  |
| --- | --- | --- | --- | --- | --- | --- | --- | --- | --- | --- | --- | --- | --- | --- | --- | --- | --- |
|  | [1] | [2] |  | [3] | [4] |  | [5] | [6] |  | [7] | [8] |  | [9] | [10] |  | [11] | [12] |
|  | Base Model | |  | Add SES | |  | Add Health | |  | Base Model | |  | Add SES | |  | Add Health | |
|  | OLS - 6% | Logit - 6% |  | OLS - 6% | Logit - 6% |  | OLS - 6% | Logit - 6% |  | OLS - 10% | Logit - 10% |  | OLS - 10% | Logit - 10% |  | OLS - 10% | Logit - 10% |
| Free T3 | 0.01 | 1.01 |  | 0.01 | 1.03 |  | 0.00 | 1.00 |  | 0.01 | 1.00 |  | 0.01 | 1.02 |  | 0.00 | 0.99 |
|  | (0.01) | (0.07) |  | (0.01) | (0.08) |  | (0.01) | (0.08) |  | (0.01) | (0.07) |  | (0.01) | (0.07) |  | (0.01) | (0.07) |
| T3 * Age | 0.04*** | 1.22*** |  | 0.02*** | 1.14*** |  | 0.01** | 1.09* |  | 0.03*** | 1.21*** |  | 0.02*** | 1.13** |  | 0.01** | 1.09* |
|  | (0.01) | (0.05) |  | (0.01) | (0.05) |  | (0.01) | (0.05) |  | (0.01) | (0.05) |  | (0.01) | (0.05) |  | (0.01) | (0.05) |
| Age (10 years) | -0.13*** | 0.45*** |  | -0.23** | 0.41 |  | -0.18* | 0.44 |  | -0.13*** | 0.46*** |  | -0.24** | 0.40 |  | -0.19* | 0.42 |
|  | (0.01) | (0.04) |  | (0.09) | (0.28) |  | (0.10) | (0.33) |  | (0.01) | (0.04) |  | (0.09) | (0.28) |  | (0.10) | (0.32) |
| Models utilize weights to account for the population sampling probabilities of the NHANES, and use linearized standard errors. | | | | | | | | | | |  |  |  |  |  |  |  |
| Coefficients come from a fully interacted model with age and the described covariates. Standard errors in parentheses. | | | | | | | | | | |  |  |  |  |  |  |  |
| T3, T4, and TSH outcomes expressed as standard deviations | | | | |  |  |  |  |  |  |  |  |  |  |  |  |  |
| Robustness of estimates to alternative samples with different levels of winsorization tested, 6% (top and bottom 3%) and 10% (top and bottom 10%). | | | | | | | | | | | | | |  |  |  |  |
| *** p<0.01, ** p<0.05, * p<0.1 | | |  |  |  |  |  |  |  |  |  |  |  |  |  |  |  |

| Table S12. Robustness of Demographic, Socio-economic, and Health Relationships to Dropping Hypo/Hyper-thyroid adults | | | | | | | |  |  |  |  |
| --- | --- | --- | --- | --- | --- | --- | --- | --- | --- | --- | --- |
|  | [1] | [2] | [3] |  | [4] | [5] | [6] |  | [7] | [8] | [9] |
| Model Specification: | Base | | |  | Base & SES | | |  | Base, SES, Health and Health Behavior | | |
|  | Free T3 | Free T4 | TSH |  | Free T3 | Free T4 | TSH |  | Free T3 | Free T4 | TSH |
| (1) Male | 0.51*** | 0.07*** | 0.01 |  | 0.52*** | 0.07** | 0.02 |  | 0.57*** | 0.06 | 0.02 |
|  | (0.02) | (0.02) | (0.02) |  | (0.02) | (0.03) | (0.02) |  | (0.04) | (0.04) | (0.04) |
| (1) Black | -0.12*** | -0.04 | -0.28*** |  | -0.16*** | -0.04 | -0.28*** |  | -0.19*** | -0.08* | -0.29*** |
|  | (0.04) | (0.04) | (0.02) |  | (0.03) | (0.04) | (0.02) |  | (0.04) | (0.05) | (0.02) |
| (1) Hispanic | 0.18*** | 0.10** | -0.10*** |  | 0.12** | 0.05 | -0.07** |  | 0.07 | 0.07 | -0.08*** |
|  | (0.04) | (0.04) | (0.02) |  | (0.05) | (0.05) | (0.03) |  | (0.05) | (0.05) | (0.03) |
| (1) Race non-white, Black, or Hispanic | -0.03 | 0.33*** | -0.15*** |  | -0.04 | 0.26*** | -0.11*** |  | 0.02 | 0.24*** | -0.10** |
|  | (0.06) | (0.06) | (0.04) |  | (0.07) | (0.05) | (0.04) |  | (0.08) | (0.06) | (0.04) |
| (1) Summer measurement | -0.10** | -0.02 | -0.00 |  | -0.11** | -0.01 | -0.00 |  | -0.12** | -0.01 | 0.01 |
|  | (0.04) | (0.06) | (0.02) |  | (0.04) | (0.06) | (0.02) |  | (0.04) | (0.06) | (0.02) |
| ln(Real HH Income) |  |  |  |  | -0.05*** | -0.02 | -0.01 |  | -0.04** | -0.02 | 0.00 |
|  |  |  |  |  | (0.02) | (0.02) | (0.01) |  | (0.02) | (0.02) | (0.01) |
| (1) College Ed. |  |  |  |  | -0.09** | 0.09*** | -0.02 |  | -0.04 | 0.09** | -0.01 |
|  |  |  |  |  | (0.04) | (0.03) | (0.03) |  | (0.04) | (0.03) | (0.03) |
| Constant | 0.33*** | -0.14* | -0.13*** |  | 0.90*** | 0.04 | -0.03 |  | 0.80*** | 0.09 | -0.11 |
|  | (0.06) | (0.07) | (0.03) |  | (0.19) | (0.22) | (0.14) |  | (0.21) | (0.22) | (0.16) |
| Observations | 7,521 | 7,521 | 7,521 |  | 7,521 | 7,521 | 7,521 |  | 6,926 | 6,926 | 6,926 |
| R-squared | 0.237 | 0.093 | 0.048 |  | 0.240 | 0.096 | 0.049 |  | 0.255 | 0.106 | 0.053 |
| Standard errors in parentheses. Models utilize weights to account for the population sampling probabilities of the NHANES, and use linearized standard errors. | | | | | | | | | | | |
| All models conditional on age, medication use, smoking, and wave. SES models conditional on household size and nativity. Health models conditional on Height, waist circumference, hours of sleep, iodine levels, calories per day, grams of sugar per day, and %HbA1c. | | | | | | | | | | | |
| Dropping individuals with hyperthyroidism (TSH<0.4mIU/L) or hypothyroidism (TSH>4.1mIU/L) | | | | | |  |  |  |  |  |  |

| Supplement Table S13. Relationships between Thyroid Hormones and Mortality, Age 50+ | | | | | | | | |
| --- | --- | --- | --- | --- | --- | --- | --- | --- |
|  | [1] | [2] | [3] | [4] |  | [5] |  | [6] |
|  | Base Model | | | |  | Add SES |  | Add Health |
|  | Mortality | | | | | | | |
| Free T3 | 0.85*** |  |  | 0.84*** |  | 0.84*** |  | 0.88** |
|  | (0.05) |  |  | (0.04) |  | (0.05) |  | (0.06) |
| Free T4 |  | 1.21*** |  | 1.23*** |  | 1.24*** |  | 1.23*** |
|  |  | (0.05) |  | (0.05) |  | (0.05) |  | (0.05) |
| TSH |  |  | 1.04 | 1.07 |  | 1.07 |  | 1.05 |
|  |  |  | (0.05) | (0.05) |  | (0.05) |  | (0.06) |
| Models utilize weights to account for the population sampling probabilities of the NHANES, and use linearized standard errors. | | | | | | | | |
| Hazard Ratios come from a Cox proportional hazards model conditional on the described covariates. Standard errors in parentheses. | | | | | | | | |
| Sample restricted to over 50 years old to focus on mortality. Mortality data through 2019. | | | | | | | | |
| Dropping individuals with hyperthyroidism (TSH<0.4mIU/L) or hypothyroidism (TSH>4.1mIU/L) | | | | | | | | |
| T3, T4, and TSH outcomes expressed as standard deviations | | | | |  |  |  |  |
| *** p<0.01, ** p<0.05, * p<0.1 | | |  |  |  |  |  |  |

| Supplement Table S14. Alternate Linear Age Specification for Demographic, Socio-economic, and Health Relationships with Standardized Thyroid-Axis Hormones | | | | | | | | | | | |  |  |  |  |
| --- | --- | --- | --- | --- | --- | --- | --- | --- | --- | --- | --- | --- | --- | --- | --- |
|  | [1] | [2] | [3] |  | [4] | [5] | [6] |  | [7] | [8] | [9] |  | [10] | [11] | [12] |
| Model Specification: | Base | | |  | Base & SES | | |  | Base, SES, Health and Health Behavior | | |  | Other Thyroid Hormones | | |
|  | Free T3 | Free T4 | TSH |  | Free T3 | Free T4 | TSH |  | Free T3 | Free T4 | TSH |  | Free T3 | Free T4 | TSH |
| Free T3 |  |  |  |  |  |  |  |  |  |  |  |  |  | 0.15*** | -0.02 |
|  |  |  |  |  |  |  |  |  |  |  |  |  |  | (0.02) | (0.01) |
| Free T4 |  |  |  |  |  |  |  |  |  |  |  |  | 0.13*** |  | -0.13*** |
|  |  |  |  |  |  |  |  |  |  |  |  |  | (0.02) |  | (0.02) |
| TSH |  |  |  |  |  |  |  |  |  |  |  |  | -0.02 | -0.12*** |  |
|  |  |  |  |  |  |  |  |  |  |  |  |  | (0.01) | (0.01) |  |
| Age in Years | -0.02*** | 0.00** | 0.01*** |  | -0.02*** | 0.00** | 0.01*** |  | -0.02*** | 0.00* | 0.01*** |  | -0.02*** | 0.01*** | 0.01*** |
|  | (0.00) | (0.00) | (0.00) |  | (0.00) | (0.00) | (0.00) |  | (0.00) | (0.00) | (0.00) |  | (0.00) | (0.00) | (0.00) |
| (1) Male | 0.51*** | 0.06** | 0.02 |  | 0.51*** | 0.06** | 0.03 |  | 0.57*** | 0.07 | 0.04 |  | 0.57*** | -0.01 | 0.06 |
|  | (0.02) | (0.03) | (0.03) |  | (0.02) | (0.03) | (0.03) |  | (0.04) | (0.04) | (0.05) |  | (0.04) | (0.04) | (0.05) |
| (1) Black | -0.12*** | -0.04 | -0.39*** |  | -0.15*** | -0.05 | -0.40*** |  | -0.18*** | -0.08* | -0.39*** |  | -0.18*** | -0.10** | -0.40*** |
|  | (0.04) | (0.04) | (0.03) |  | (0.04) | (0.04) | (0.04) |  | (0.04) | (0.04) | (0.04) |  | (0.04) | (0.05) | (0.04) |
| (1) Hispanic | 0.18*** | 0.10** | -0.12*** |  | 0.12** | 0.04 | -0.09* |  | 0.07 | 0.06 | -0.10** |  | 0.06 | 0.03 | -0.09* |
|  | (0.03) | (0.04) | (0.03) |  | (0.05) | (0.05) | (0.05) |  | (0.05) | (0.06) | (0.05) |  | (0.05) | (0.06) | (0.05) |
| (1) Race non-white, Black, or Hispanic | -0.01 | 0.32*** | -0.16*** |  | -0.02 | 0.25*** | -0.12** |  | 0.04 | 0.22*** | -0.09 |  | 0.01 | 0.20*** | -0.06 |
|  | (0.06) | (0.06) | (0.04) |  | (0.07) | (0.05) | (0.05) |  | (0.08) | (0.06) | (0.06) |  | (0.08) | (0.06) | (0.06) |
| (1) Summer measurement | -0.09* | -0.03 | -0.00 |  | -0.09** | -0.02 | -0.00 |  | -0.11** | -0.02 | 0.00 |  | -0.10** | -0.01 | -0.00 |
|  | (0.04) | (0.06) | (0.03) |  | (0.04) | (0.06) | (0.03) |  | (0.05) | (0.06) | (0.03) |  | (0.05) | (0.06) | (0.03) |
| ln(Real HH Income) |  |  |  |  | -0.04** | -0.04** | -0.04** |  | -0.03* | -0.04** | -0.01 |  | -0.03 | -0.04** | -0.02 |
|  |  |  |  |  | (0.02) | (0.02) | (0.01) |  | (0.02) | (0.02) | (0.02) |  | (0.02) | (0.02) | (0.02) |
| (1) College Ed. |  |  |  |  | -0.10*** | 0.07** | 0.01 |  | -0.04 | 0.05 | 0.04 |  | -0.05 | 0.06** | 0.04 |
|  |  |  |  |  | (0.03) | (0.03) | (0.04) |  | (0.04) | (0.03) | (0.05) |  | (0.04) | (0.03) | (0.05) |
| Constant | -0.21*** | -0.21*** | 0.20*** |  | 0.29 | 0.21 | 0.59*** |  | 0.19 | 0.22 | 0.35* |  | 0.17 | 0.23 | 0.38** |
|  | (0.05) | (0.06) | (0.04) |  | (0.18) | (0.21) | (0.16) |  | (0.20) | (0.21) | (0.18) |  | (0.20) | (0.21) | (0.18) |
| Observations | 8,059 | 8,059 | 8,059 |  | 8,059 | 8,059 | 8,059 |  | 7,426 | 7,426 | 7,426 |  | 7,426 | 7,426 | 7,426 |
| R-squared | 0.241 | 0.080 | 0.051 |  | 0.245 | 0.082 | 0.052 |  | 0.260 | 0.090 | 0.059 |  | 0.276 | 0.124 | 0.075 |
| Standard errors in parentheses. Models utilize weights to account for the population sampling probabilities of the NHANES, and use linearized standard errors. | | | | | | | | | | | |  |  |  |  |
| All models conditional on age, medication use, smoking, and wave. SES models conditional on household size and nativity. Health models conditional on Height, waist circumference, hours of sleep, iodine levels, calories per day, grams of sugar per day, and %HbA1c. | | | | | | | | | | | | | | | |
| T3, T4, and TSH outcomes expressed as standard deviations | | |  |  |  |  |  |  |  |  |  |  |  |  |  |
| *** p<0.01, ** p<0.05, * p<0.1 |  |  |  |  |  |  |  |  |  |  |  |  |  |  |  |
